# Oct4 primarily controls enhancer activity rather than accessibility

**DOI:** 10.1101/2021.06.28.450119

**Authors:** Le Xiong, Erik A. Tolen, Jinmi Choi, Livia Caizzi, Kenjiro Adachi, Michael Lidschreiber, Patrick Cramer, Hans R. Schöler

## Abstract

The transcription factor Oct4 is essential for maintaining stem cell pluripotency and for efficient cell reprogramming, but its functional roles are far from being understood. Here, we investigate the functions of Oct4 by rapidly depleting Oct4 from mouse embryonic stem cells and conducting a time-resolved multiomics analysis. Oct4 depletion leads to an immediate loss of its binding to putative enhancers that are accessible in chromatin. Loss of Oct4 is accompanied by a concomitant decrease in mRNA synthesis from putative target genes that are part of the transcriptional network that maintains pluripotency. Oct4 binding to enhancers does not correlate with chromatin accessibility, whereas Sox2 can apparently retain accessibility after Oct4 depletion even in the absence of eRNA synthesis. These results are consistent with the model that Sox2 primarily acts as a pioneer factor that renders enhancers accessible, whereas Oct4 acts primarily as a transcriptional activator that stimulates transcription of pluripotency enhancers and their target genes.

## Introduction

The transcription factor (TF) Oct4 is essential for maintaining pluripotency *in vitro* (Jerabek et al., 2014) as well as *in vivo* (Nichols et al., 1998). Oct4 lies at the core of an intricate transcriptional regulatory network that maintains the pluripotent state (Boyer et al., 2005; Zhou et al., 2007). Oct4 binds to enhancers (Schöler et al., 1989), which are cis-regulatory genomic elements that orchestrate gene expression in metazoan (Banerji et al., 1981). Oct4 function in maintaining of embryonic stem cells (ESCs) pluripotency has been attributed to the establishment of super-enhancers (SEs), which show high occupancy with TFs and coactivators (Hnisz et al., 2013; Whyte et al., 2013). In mouse ESCs, the loss of Oct4 decreases the activity of pluripotency genes that are located near SEs (Whyte et al., 2013). Degradation of Oct4 leads to preferential decrease of Oct4 and Mediator occupancy at SEs (Boija et al., 2018).

Oct4 cooperates with the TF Sox2 at thousands of genomic sites in ESCs (Ambrosetti et al., 1997; Chen et al., 2008; Loh et al., 2006; Yuan et al., 1995). Oct4 and Sox2 can bind enhancers adjacently to well-defined composite DNA motifs (Chen et al., 2008; Lam et al., 2012). Two recent studies showed that depletion of Oct4 from ESCs for 24 hours resulted in a loss of chromatin accessibility at a majority of its occupied sites, accompanied by a loss of Sox2 binding (Friman et al., 2019; King and Klose, 2017). This suggested a role of Oct4 in the control of chromatin accessibility (Friman et al., 2019; King and Klose, 2017). However, secondary effects could not be excluded, and thus the contributions of Oct4 and Sox2 to chromatin accessibility and enhancer function remain unclear.

Enhancers are often transcribed to produce enhancer RNA (eRNA) (De Santa et al., 2010; Kim et al., 2010; Tuan et al., 1992). The functions of enhancer transcription and eRNA remain poorly understood (Lewis et al., 2019; Li et al., 2016; Sartorelli and Lauberth, 2020). Transcribing Pol II at enhancers contributes to chromatin alterations by recruiting histone modifying and remodeling factors (Ho et al., 2006; Kaikkonen et al., 2013; Ling et al., 2004), and eRNA may be involved in gene regulation (Bose et al., 2017; Gorbovytska et al., 2021; Kaikkonen et al., 2013; Li et al., 2013; Mousavi et al., 2013; Schaukowitch et al., 2014; Sigova et al., 2015). The synthesis of eRNA correlates with the transactivation activity of enhancers (Andersson et al., 2014; De Santa et al., 2010; Djebali et al., 2012; Hah et al., 2013; Henriques et al., 2018; Kim et al., 2010; Li et al., 2013; Lidschreiber et al., 2021; Melgar et al., 2011; Michel et al., 2017; Schwalb et al., 2016; Wu et al., 2014). The synthesis of eRNA can be used to identify putative enhancers by transient transcriptome sequencing (TT-seq), a method that captures newly synthesized RNA (Schwalb et al., 2016). TT-seq combines a short pulse of 4-thiouridine (4sU) labeling with RNA fragmentation and monitors transcription changes at both enhancers and their target genes genome-wide (Schwalb et al., 2016). TT-seq can quantify changes in enhancer transcription during cellular processes such as T-cell stimulation, the heat shock response or transdifferentiation, and is thus ideal to monitor immediate transcriptome changes after perturbation (Choi et al., 2021; Gressel et al., 2019; Michel et al., 2017).

To investigate the primary function of Oct4 in the control of pluripotency, we used rapid depletion of Oct4 in mouse ESCs. We then conducted a time course study to monitor changes in the transcriptome by TT-seq, changes in chromatin accessibility by ATAC-seq (Buenrostro et al., 2013), and changes in Oct4 and Sox2 occupancy by ChIP-seq. We found that Oct4 depletion leads to a rapid loss of eRNA synthesis and Oct4 binding to enhancers, which correlated with a decrease in mRNA synthesis from nearby putative target genes. In contrast, chromatin accessibility at Oct4-bound enhancers was generally decreased only later, arguing that Oct4 primarily functions in transcription activation, not chromatin opening, which depends on Sox2. Taken together, these results suggest that Oct4 maintains pluripotency in ESCs primarily by controlling the activity, rather than the accessibility, of enhancers.

## Results

### Rapid Oct4 depletion and transcription unit annotation

To investigate the role of Oct4 in maintaining pluripotency, we used a doxycycline (DOX) inducible Oct4 loss-of-function mouse embryonic stem cell line (mESC) ZHBTc4 that allows for rapid depletion of Oct4 (Niwa et al., 2000). This system was previously used to study the effect of Oct4 depletion after 24 hours of DOX treatment (Friman et al., 2019; King and Klose, 2017). In order to investigate the direct, primary role of Oct4, we conducted a time course experiment collecting samples after 0, 3, 6, 9, 12 and 15 hours of DOX treatment. We found that Oct4 protein levels were already reduced after 6 hours of DOX treatment and decreased strongly after 9 hours (**Figure 1A**, **whole cell lysate**). Oct4 protein depletion was nearly complete at 24 hours of treatment, while Sox2 and Nanog protein levels remained essentially unchanged for extended times before decreasing (**Figure 1A, Figure 1-figure supplement 1A**). Chromatin binding of Oct4 was impaired already after 3 hours of DOX treatment, whereas Sox2 binding decreased after 9 hours and Nanog binding remained unchanged over the entire time course (**Figure 1A, chromatin**).

**Figure 1.**
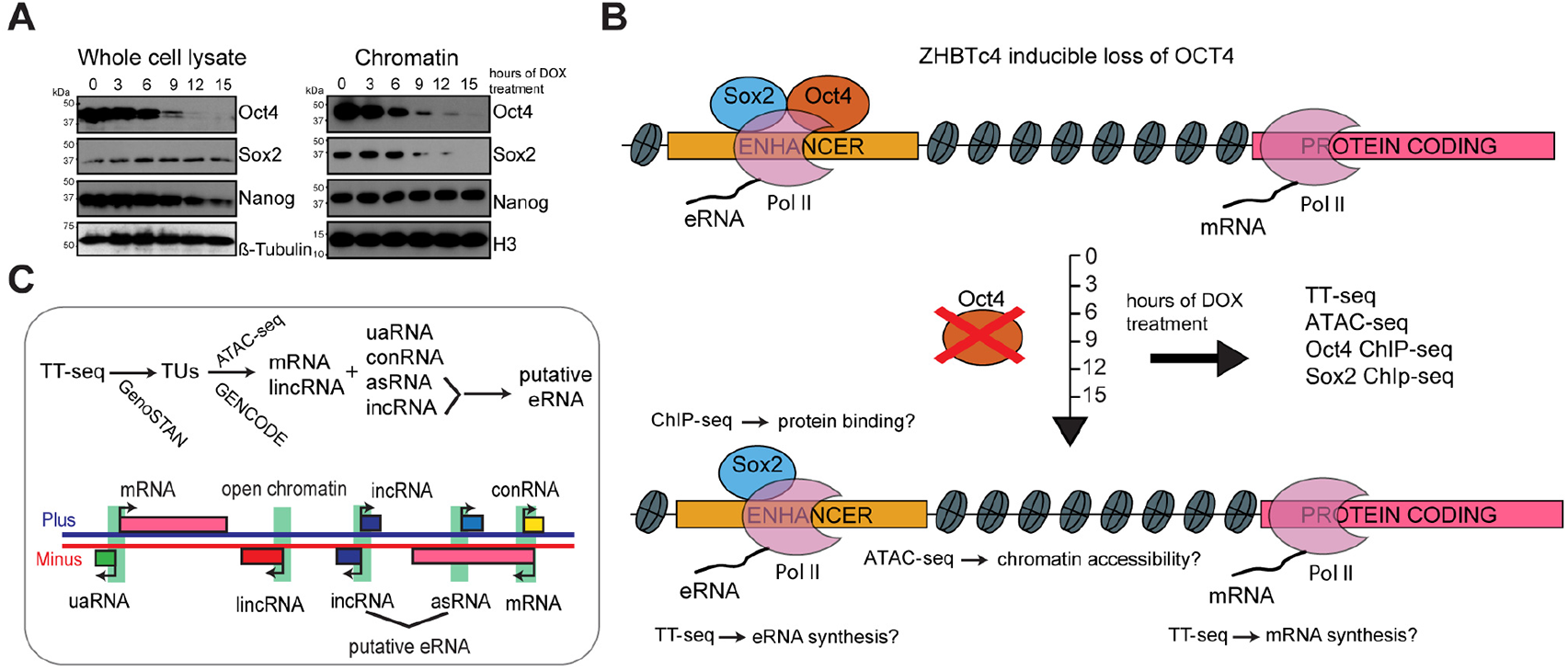
Rapid Oct4 depletion in ZHBTc4 mouse ESCs. **(A)** Western blot analysis of whole cell lysate and chromatin samples over time course of DOX treatment using Oct4, Sox2, Nanog, β-Tubulin (control) and H3 (control) antibodies. **(B)** Schematic of methodology and samples collected. TT-seq, ATAC-seq, Oct4 and Sox2 ChIP-seq experiments were performed after 0, 3, 6, 9, 12 and 15 hours of DOX treatment in ZHBTc4 mouse ESCs. **(C)** Transcription unit annotation. Genome segmentation with GenoSTAN was used to annotate transcription units (TUs) from TT-seq data. ATAC-seq data and mouse GENCODE annotation were then used to classify TUs.

To monitor the effect of a rapid Oct4 depletion on transcription, we conducted TT-seq (Schwalb et al., 2016) at 0, 3, 6, 9, 12 and 15 hours after DOX treatment (**Figure 1B**). RNA labelling with 4-thiouridine (4sU) was carried out for 5 minutes (min) and two independent biological replicates were generated for each time point (**Table S1**). To study the role of Oct4 in maintaining chromatin accessibility, we performed ATAC-seq (Buenrostro et al., 2013) over the same time course (**Figure 1B, Table S2**). TT-seq and ATAC-seq data were highly reproducible (**Figure 1-figure supplement 1B-C**).

We then used the TT-seq data to segment the genome into transcription units (TUs) and non-transcribed regions using GenoSTAN (Zacher et al., 2017) (**Figure 1C and Methods**). To avoid spurious predictions, TUs detected by TT-seq had to exceed a minimal expression cutoff of RPK > 26.5 and had to originate from an open chromatin region identified by ATAC-seq (**Figure 1-figure supplement 1D-E**). We sorted TUs into protein-coding RNAs (mRNAs) and long intergenic non-coding RNAs (lincRNAs) based on GENCODE annotation (Frankish et al., 2019). The remaining non-coding TUs were classified as upstream antisense RNA (uaRNA), convergent RNA (conRNA), antisense RNA (asRNA) and intergenic RNA (incRNA) units according to the location relative to mRNA (**Figure 1C and Methods**). This resulted in 9,266 mRNAs, 9,257 incRNAs, 3,661 asRNAs, 1,981 uaRNAs, 471 conRNAs and 318 lincRNAs (**Figure 1-figure supplement 1F**). The length distribution of the detected RNA units (**Figure 1-figure supplement 1G**) agreed with previous estimations (Michel et al., 2017; Schwalb et al., 2016).

### Oct4 maintains the transcriptional network governing pluripotency

We first investigated changes in mRNA synthesis during Oct4 depletion. Changes could already be observed after 3 hours of DOX treatment (**Figure 2-figure supplement 1A**), in agreement with chromatin fractionation results (**Figure 1A, chromatin**). Differential gene expression analysis (Love et al., 2014) detected 769 significantly down-regulated and 829 up-regulated genes (adjusted P-value = 0.01) after 15 hours of DOX treatment (**Figure 2A-B, Figure 2-figure supplement 1B**). To dissect the kinetics of mRNA synthesis changes of differentially expressed genes, we performed k-means clustering and classified early and late down- and up-regulated genes (**Table S3**). Early down-regulated genes (446 genes) showed the strongest decrease in mRNA synthesis after 6-9 hours of DOX treatment (**Figure 2C-D, Figure 2-figure supplement 1C, left**), whereas late down-regulated genes (323 genes) decreased most strongly after 12-15 hours (**Figure 2C-D, Figure 2-figure supplement 1C, right**). Early and late up-regulated genes behaved similarly (**Figure 2-figure supplement 1D-F**). Gene ontology (GO) analysis (Huang da et al., 2009) showed that early down-regulated genes were enriched for stem cell population maintenance (**Figure 2E**), whereas late down-regulated genes were enriched for DNA replication and cell cycle (**Figure 2-figure supplement 1G**). Early up-regulated genes showed enrichment for carbohydrate metabolic process (**Figure 2-figure supplement 1H**), whereas late up-regulated genes were slightly enriched for *in utero* embryonic development (**Figure 2-figure supplement 1I**). These findings reflect the known transition of ZHBTc4 cells towards the trophectoderm upon loss of Oct4 (Niwa et al., 2000).

**Figure 2.**
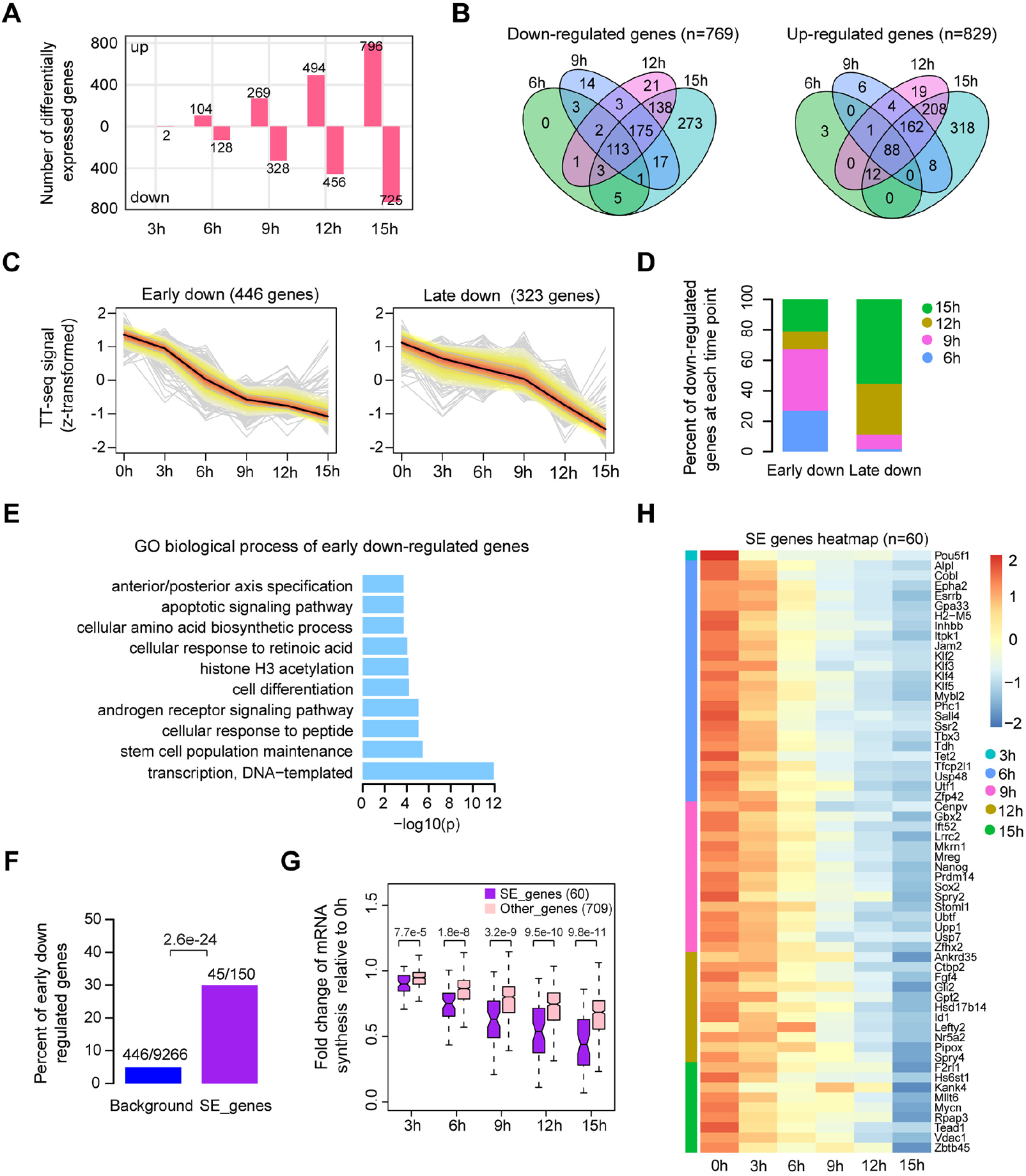
Oct4 maintains the transcriptional network governing pluripotency. **(A)** Number of differentially expressed mRNAs detected by DESeq2 after each time point of DOX treatment. **(B)** Venn diagram showing the overlap for differentially expressed mRNAs detected by DESeq2 at each time point of DOX treatment. **(C)** K-means clustering of 769 down-regulated mRNAs into early (left) and late (right) down-regulated groups. Y axis indicates z-score transformed TT-seq counts. **(D)** Differentially regulated time point composition for early (left) and late (right) down-regulated gene groups. Y axis indicates percentage. **(E)** GO biological process enrichment of early down-regulated mRNAs. **(F)** Bar graph depicting the percentage of early down-regulated mRNAs in all annotated mRNAs (blue, as background) and SE-controlled mRNAs (purple). SE annotation was obtained from (Whyte et al., 2013). P-value was calculated by Fisher’s exact test. **(G)** Boxplot indicating the changes in mRNA synthesis for putative SE-controlled down-regulated genes (n=60, purple) versus other down-regulated genes (n=709, pink). Y axis indicates fold change of mRNA synthesis relative to 0 hours. P-values were calculated by Wilcoxon rank sum test. Black bars represent the median values for each group. Lower and upper boxes are the first and third quartiles, respectively. The ends of the whiskers extend the box by 1.5 times the interquartile range. Outliers are not drawn. **(H)** Heatmap indicating the kinetics of SE-controlled down-regulated genes (n=60). Genes were ordered by the corresponding time of significant down regulation.

We then assessed if there is an enrichment of putative SE-controlled genes among the early down-regulated genes. Indeed, of the 150 transcribed genes that are nearest to SEs (Whyte et al., 2013), we found 60 genes to be significantly down-regulated, of which 45 were early down-regulated (**Figure 2F**, **P-value=2.6e-24, Fisher’s exact test**). We then compared the kinetics of mRNA synthesis changes of the putatively SE-controlled down-regulated genes (60) to other down-regulated genes (709). This showed that mRNA synthesis of SE-controlled down-regulated genes was particularly sensitive to Oct4 depletion (**Figure 2G**). Among the 60 SE genes that were down-regulated, we found many pluripotency genes at early time points (**Figure 2H**). At 6 hours of Oct4 depletion we found a significant decrease of synthesis for *Esrrb*, *Klf2*, *Klf4*, *Utf1* and *Tbx3*. At 9 hours of depletion, *Sox2*, *Nanog* and *Prdm14* were significantly down-regulated, and *Nr5a2* and *Fgf4* after 12 hours. Taken together, our analysis of early mRNA synthesis changes upon Oct4 depletion revealed a rapid downregulation of the components of pluripotency transcriptional network with SE-controlled genes being immediately and strongly affected. Thus, consistent with previous findings (Whyte et al., 2013), Oct4 is strictly required to maintain the transcriptional network underlying pluripotency.

### Oct4-bound transcribed enhancers produce high levels of eRNA

To understand how loss of Oct4 leads to rapid destabilization of the pluripotency gene network, we combined our TT-seq and ATAC-seq data with published Oct4 ChIP-seq data (King and Klose, 2017) to derive a refined annotation of putative enhancers in mESCs (**Figure 3A**). First, we defined transcribed enhancers by annotating putative eRNAs (**Figure 3A**). We selected asRNAs and incRNAs that originated over 1 kb away from promoter-related RNAs (mRNA, conRNA and uaRNA) and merged those located less than 1 kb apart (**Figure 1C**). This resulted in 8,727 putative eRNAs, consisting of 2,468 asRNAs and 6,259 incRNAs, with a median length ∼700 bp (**Figure 3-figure supplement 1A-B**). To annotate putative Oct4-regulated eRNAs, we used available Oct4 ChIP-seq data (King and Klose, 2017) (**Table S4**). Most Oct4 ChIP-seq peaks (91%) overlapped with open chromatin regions identified by ATAC-seq (**Figure 3-figure supplement 1C**). Out of the 8,727 putative eRNAs, 2,221 overlapped with 2,231 Oct4-bound sites (Klf4 SE shown as an example in **Figure 3-figure supplement 1D**). We refer to these Oct4-bound sites as Oct4-bound transcribed enhancers (**Figure 3A**). The majority of Oct4-bound transcribed enhancers (90%) were marked by histone H3 lysine 4 mono-methylation (H3K4me1) (**Figure 3-figure supplement 1E**). Oct4-bound transcribed enhancers were strongly enriched for biological processes related to stem cell population maintenance (**Figure 3-figure supplement 1F**). In contrast, the Oct4-unbound transcribed enhancers (**Figure 3A**) were enriched for other biological processes (**Figure 3-figure supplement 1G**). Moreover, Oct4-regulated eRNAs originating from Oct4-bound enhancers were significantly longer and showed higher synthesis than other putative eRNAs (**Figure 3-figure supplement 1E**, P-value < 2.2e-16, Wilcoxon rank sum test).

**Figure 3.**
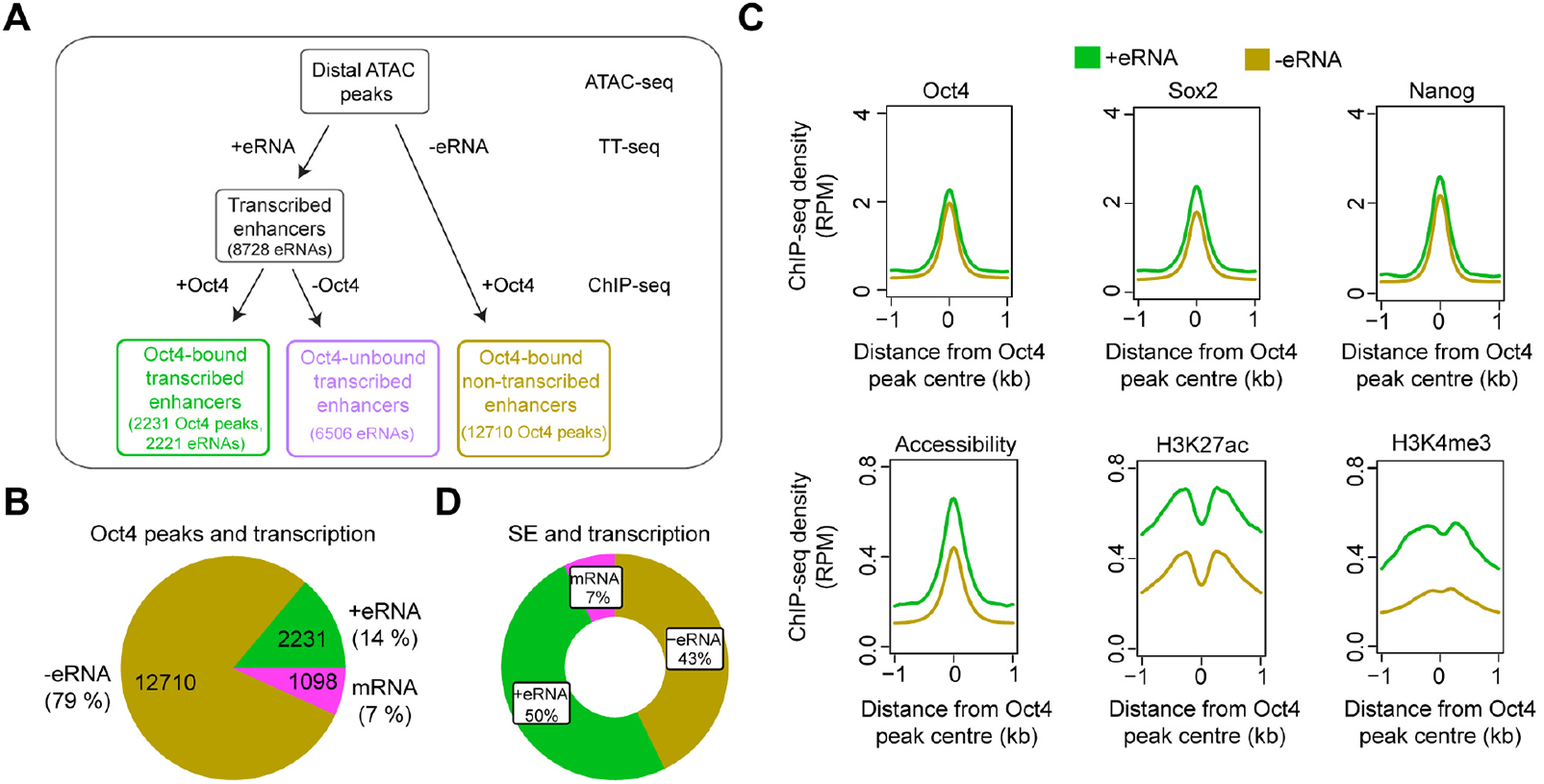
Annotation of putative Oct4-bound/regulated enhancer classes in mESCs. **(A)** Diagram illustrating classification of Oct4 binding sites by combing ATAC-seq, TT-seq annotated eRNAs, and Oct4 ChIP-seq peaks. Oct4 ChIP-seq data at 0 hours DOX treatment was obtained from (King and Klose, 2017). **(B)** Pie chart indicating the overlap of Oct4 binding sites with regions of active transcription (eRNA or mRNA) or no transcription annotated by TT-seq. **(C)** Metagene plot showing the occupancy for transcription factors Oct4, Sox2, Nanog, chromatin accessibility, and H3K27ac and H3K4me3 histone modifications at Oct4-bound transcribed enhancers (n=2,231) and Oct4-bound non-transcribed enhancers (n=12,710). Y axis depicts ChIP-seq coverage density in reads per million (RPM). ChIP-seq data of Oct4, Sox2 and Nanog were obtained from (King and Klose, 2017), H3K27ac and H3K4me3 histone modifications data were obtained from (Chronis et al., 2017). **(D)** Pie chart depicting the overlap of Oct4-bound sites (n=514) at SE that show eRNA transcription (n=256, 50%), mRNA transcription (n=38, 7%) or no transcription (n=220, 43%) by TT-seq.

Of the remaining Oct4-bound accessible sites, 1,098 produced mRNAs, and 12,710 produced no detectable RNAs and were referred to as Oct4-bound non-transcribed enhancers (**Figure 3A, B**). We then performed metagene analysis for TF enrichment at Oct4-bound transcribed and non-transcribed enhancers using published data (King and Klose, 2017; Chronis et al., 2017) (**Table S4**). Whereas both groups of enhancers showed similar H3K4me1 levels, Oct4-bound transcribed enhancers showed an enrichment with active histone marks H3K27ac and H3K4me3, higher chromatin accessibility, and higher occupancies with Oct4, Sox2, Nanog, Klf4 and Esrrb (**Figure 3C, Figure 3-figure supplement 1H**). Moreover, Oct4-bound transcribed enhancers were located closer to their nearest active putative target genes (median distance of 37 kb) as compared to non-transcribed enhancers (median distance 89 kb) (**Figure 3-figure supplement 1I**). Finally, we investigated eRNA synthesis at SEs. Half of Oct4-bound sites within SEs produced eRNAs (**Figure 3D**). The eRNAs obtained from SEs were generally longer and had higher synthesis compared to eRNAs from typical enhancers (TEs), and SEs showed higher occupancy with Oct4, Sox2, Nanog, Klf4 and Esrrb (**Figure 3-figure supplement 1J-K**). These efforts led to an enhancer annotation in mESCs and suggested that Oct4-bound enhancers are transcriptionally more active than other enhancers.

### Oct4 is often required for enhancer transcription

We next analyzed Oct4-bound transcribed enhancers (**Figure 3A**) with respect to changes in their eRNA synthesis upon Oct4 depletion. Synthesis of eRNAs was highly reproducible between the two biological replicates (**Figure 4-figure supplement 1A**). PCA analysis revealed that the changes of Oct4-regulated eRNA synthesis followed a similar trajectory to that seen for mRNAs (**Figure 4-figure supplement 1B, Figure 2-figure supplement 1A**). Differential expression analysis of eRNAs (Love et al., 2014) detected significant down-regulation of 782 Oct4-regulated eRNAs after 15 hours of DOX treatment (**Figure 4A, Figure 4-figure supplement 1C, adjusted P-value = 0.01**). The kinetics analysis showed that for the down-regulated eRNAs synthesis decreased already after 3 hours (**Figure 4B**). Moreover, SE eRNAs were more strongly down-regulated compared to TE eRNAs (**Figure 4C-4D**). Taken together, these results suggest that Oct4 is required for eRNA synthesis at about one third of putative Oct4-bound transcribed enhancers including SEs.

**Figure 4.**
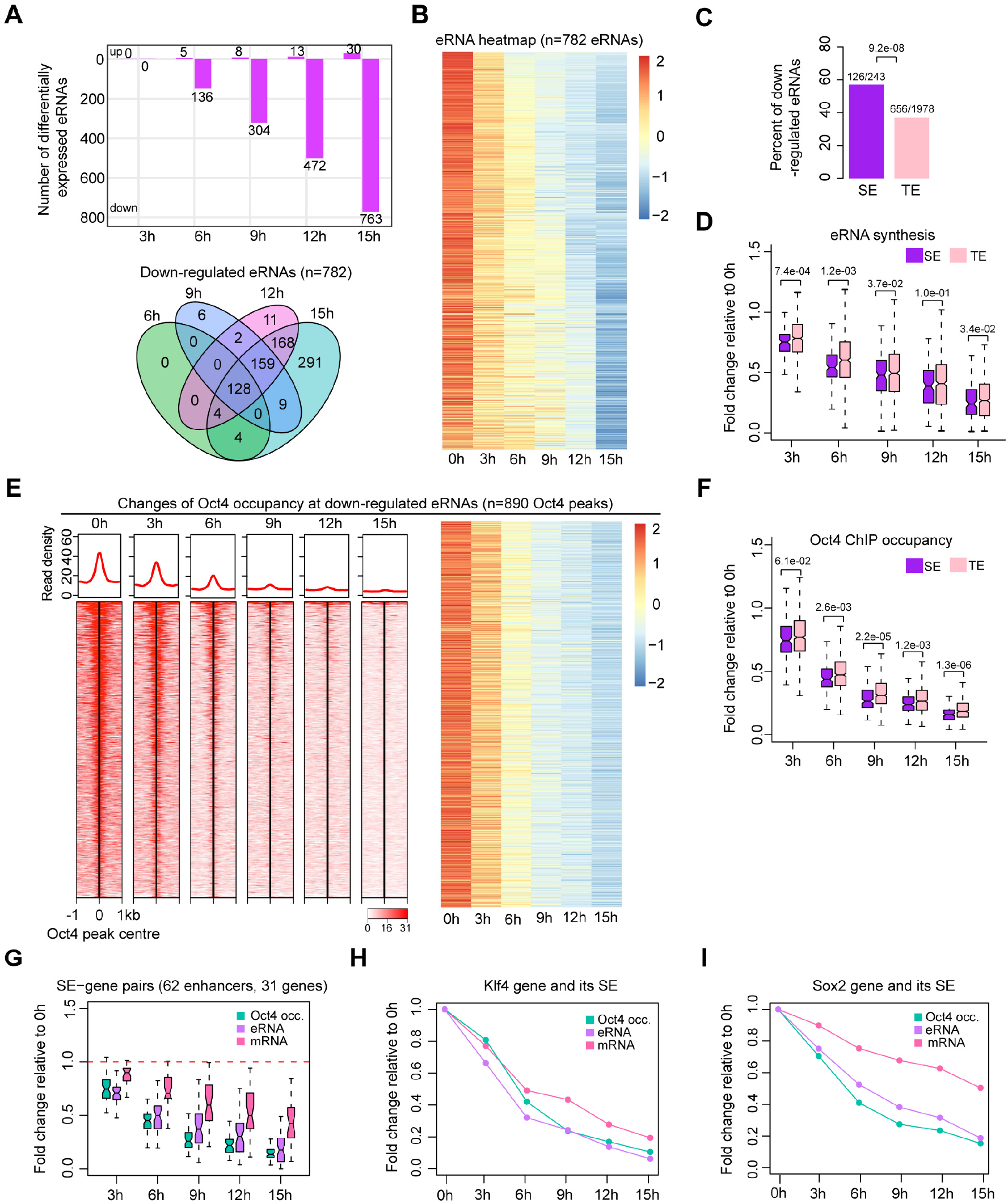
Oct4 is required for enhancer and gene transcription. **(A)** Number of differentially expressed Oct4-regulated eRNAs detected by DESeq2 (top) and Venn diagram showing overlapping of differentially expressed Oct4-regulated eRNA (bottom) at each time point of DOX treatment. **(B)** Heatmap visualizing the kinetics of Oct4-regulated down-regulated eRNAs (n=782). **(C)** Bar chart indicating the percentage of down-regulated eRNAs at SE and TE. P-value was calculated by Fisher’s exact test. **(D)** Boxplot indicating synthesis changes of down-regulated eRNAs at SE (n=126) versus TE (n=656). Y axis indicates fold change of eRNAs synthesis relative to 0 hours. P-values were calculated by Wilcoxon rank sum test. Black bars represent the median values for each group. Lower and upper boxes are the first and third quartiles, respectively. The ends of the whiskers extend the box by 1.5 times the interquartile range. Outliers are not drawn. **(E)** Changes of Oct4 occupancy at down-regulated eRNAs, illustrated by Oct4 ChIP-seq coverage (left) and count heatmaps (right). 782 down-regulated eRNAs were originated from 890 Oct4-bound transcribed enhancers (peaks). Normalized read densities were presented, in left, peaks were ranked by Oct4 read densities. **(F)** Boxplot indicating the corresponding Oct4 occupancy changes at down-regulated SE versus TE eRNAs defined in (**D**). Y axis indicates fold change of Oct4 occupancy relative to 0 hours. P-values were calculated by Wilcoxon rank sum test. **(G)** Boxplot indicating changes of Oct4 occupancy, eRNA and mRNA synthesis for 62 SE-gene pairs. Y axis indicates fold change relative to 0h. Transcriptionally down-regulated SEs were paired with their nearest transcribed genes and pairs with down-regulated genes were kept. Enhancers number were counted by individual Oct4 peaks within SEs. **(H)** Fold changes of Oct4 occupancy, eRNA and mRNA synthesis at *Klf4* gene and its associated SE. Fold changes of Oct4 occupancy and eRNA synthesis at SE were represented by average of the individual enhancers within the SE (illustrated as IGV track in Figure 5F). **(I)** Fold changes of Oct4 occupancy, eRNA and mRNA synthesis at *Sox2* gene and its associated SE. Fold changes of Oct4 occupancy and eRNA synthesis at SE were represented by average of the individual enhancers within the SE (illustrated as IGV track in Figure 5G).

### Oct4 binds enhancers to activate putative target genes

To investigate whether Oct4 depletion leads to a loss of Oct4 binding to enhancers, we performed ChIP-seq of Oct4 after 0, 3, 6, 9, 12 and 15 hours of DOX treatment (**Table S5**). In agreement with chromatin fractionation results (**Figure 1A**, **chromatin**), Oct4 occupancy decreased after 3 hours and strongly dropped after 9 hours of DOX treatment (**Figure 4E**). This is consistent with the observed decrease in eRNA synthesis (**Figure 4B**). In accordance with our previous results (**Figure 4D**), occupancy of Oct4 decreased more strongly at SEs compared to TEs (**Figure 4F**). These results show that Oct4 binding is required for eRNA synthesis at a subset of Oct4-bound transcribed enhancers and particularly at SEs.

To investigate whether the observed decrease in Oct4 occupancy and eRNA synthesis at Oct4-bound transcribed enhancers coincided with a decrease of target mRNA synthesis, we then paired transcriptionally down-regulated SEs with their nearest transcribed genes and kept those pairs with down-regulated genes. This resulted in 62 enhancer-gene pairs. For them, we found that eRNA and mRNA synthesis decreased already after 3 hours of DOX treatment and occurred with similar trajectories over the entire time course, coinciding with a decrease in Oct4 occupancy (**Figure 4G**). This is further illustrated for *Klf4* and *Sox2* genes. Synthesis of both genes decreased at 3 hours and changes over time correlated well with changes of Oct4 occupancy and eRNA synthesis at their well-studied SEs (**Figure 4H-I**). These results are consistent with a function of Oct4 in activating both enhancer transcription and mRNA synthesis from putative target genes.

### Oct4 binding does not directly correlate with enhancer accessibility

We next investigated changes in chromatin accessibility at Oct4-bound transcribed enhancers. Principal component analysis indicated that accessibility changes started to occur after 6 hours of DOX treatment, with substantial changes until 12 hours and followed by a subtle change at 15 hours (**Figure 5-figure supplement 1A**). To call significantly changed accessible chromatin regions we used DESeq2 (Love et al., 2014). In contrast to mRNAs and eRNAs (**Figure 2A, Figure 4A**), only few enhancers were detected to have significantly altered chromatin accessibility at 6 hours (**Figure 5A, Figure 5-figure supplement 1B**). 15 hours of DOX treatment resulted in a significant decrease of chromatin accessibility at 726 enhancers (adjusted P-value = 0.01) (**Figure 5A, Figure 5-figure supplement 1B**). The kinetic analysis showed that for these enhancer, chromatin accessibility remained unchanged at 3 hours (**Figure 5B**). These results show that decreased Oct4 binding does not directly correlate with chromatin accessibility changes, which occur later than Oct4 depletion. This delayed effect on chromatin accessibility argues that the primary function of Oct4 is not to render or keep chromatin accessible.

**Figure 5.**
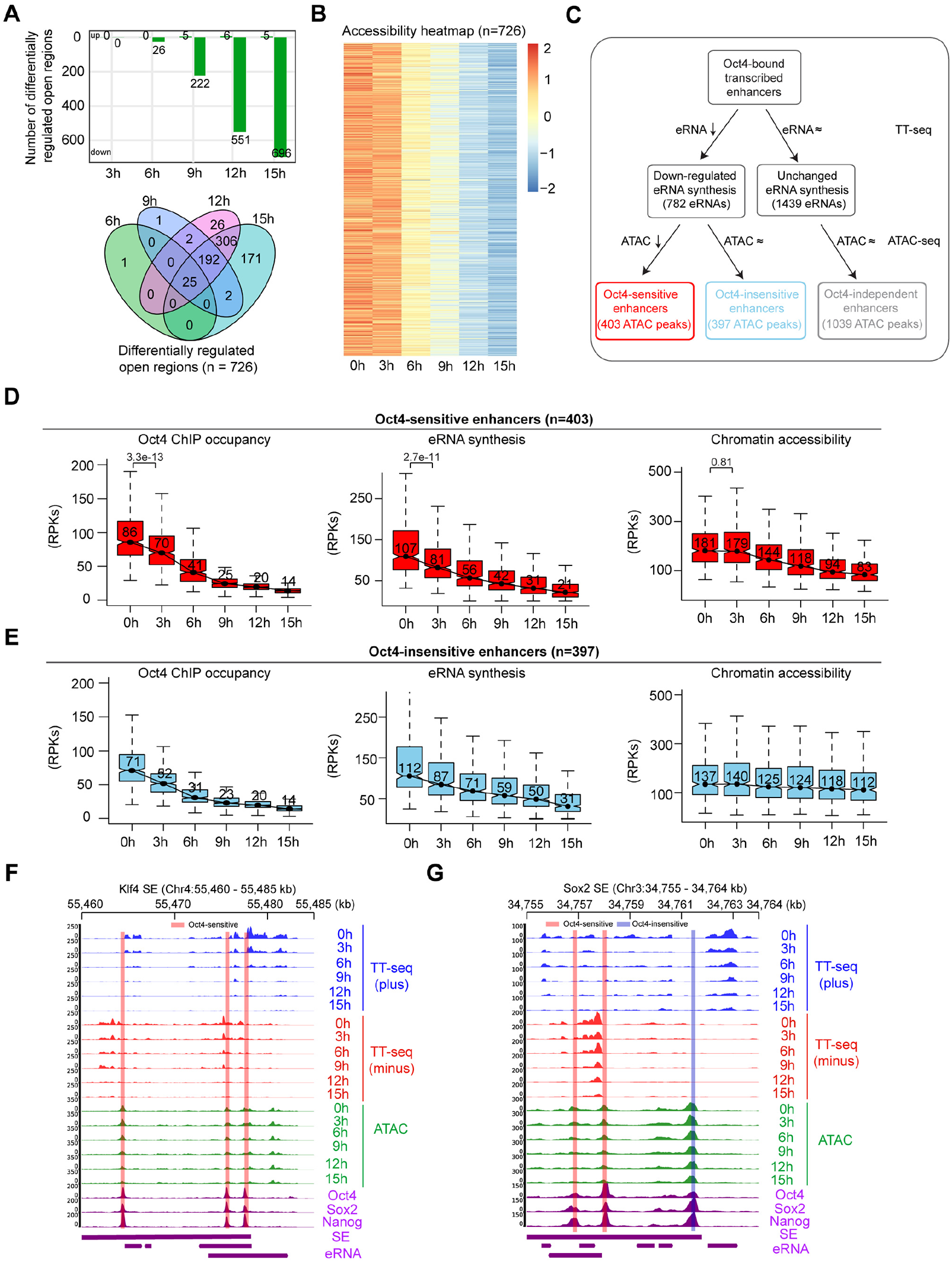
Oct4 binding does not generally correlate with enhancer accessibility. **(A)** Number of differentially regulated chromatin open regions detected by DESeq2 for Oct4-bound transcribed enhancers (top) and Venn diagram showing overlapping of detected differentially regulated open regions (bottom) for each time point of DOX treatment. **(B)** Heatmap visualizing the kinetics of chromatin accessibility changes at Oct4-occupied accessibility decreased sites (n=726). **(C)** Diagram indicating classification of Oct4-sensititive, -insensitive and -independent enhancers defined by change of eRNA synthesis and chromatin accessibility at Oct4-bound transcribed enhancers. **(D)** Boxplots indicating the changes of Oct4 occupancy, eRNA synthesis and chromatin accessibility at Oct4-sensitive enhancers (n=403). P-values were calculated by Wilcoxon rank sum test. Y axis represents read counts per kilobases (RPKs). Black bars represent the median values for each group. Lower and upper boxes are the first and third quartiles, respectively. The ends of the whiskers extend the box by 1.5 times the interquartile range. Outliers are omitted. **(E)** Boxplots indicating the changes of Oct4 occupancy, eRNA synthesis and chromatin accessibility at Oct4-insensitive enhancers (n=397). **(F)** Genome browser view for changes of eRNA synthesis and chromatin accessibility at *Klf4* SE including three Oct4-sensitive enhancers. Tracks from top to bottom: TT-seq coverages of plus strand (blue), minus strand (red) and ATAC-seq coverages (green) at 0, 3, 6, 9, 12 and 15 hours; ChIP-seq coverages of Oct4, Sox2 and Nanog (purple) from ZHBTc4 mouse ES cell at 0 hours (King and Klose, 2017); SE annotation (Whyte et al., 2013); *Klf4* SE eRNAs annotated by TT-seq (purple arrow). Biological replicates were merged for visualization. **(G)** Genome browser view for changes of eRNA synthesis and chromatin accessibility at *Sox2* SE including two Oct4-sensitive enhancers and one Oct4-insensitive enhancer. Tracks were ordered in the same way as in (**F**).

To further investigate whether Oct4 may have roles in altering chromatin, we classified the down-regulated Oct4-bound transcribed enhancers (**Figure 4B**) based on their respective changes in chromatin accessibility. We defined two groups of down-regulated transcribed enhancers, showing either decreased or unchanged chromatin accessibility (**Figure 5C**). In both groups, Oct4 binding to chromatin decreased over the time course and was associated with a decrease in eRNA synthesis (**Figure 5D-E**). For the first group, depletion of Oct4 led to a decrease in eRNA synthesis and a reduction of chromatin accessibility, with down regulation of eRNA synthesis preceding the decrease in accessibility (**Figure 5D**). This is illustrated for the SE near *Klf4* gene, which contains three enhancers of this group (**Figure 5F**). Synthesis of *Klf4* gene decreased at 3 hours and occurred simultaneously with decrease of Oct4 occupancy and eRNA synthesis at its SE (**Figure 4H**), whereas the corresponding chromatin accessibility started to decrease at 9 hours (**Figure 5F**). For the second group, depletion of Oct4 led to a decrease in eRNA synthesis without changes in chromatin accessibility (**Figure 5E**). This is illustrated by the *Sox2* gene SE and *Mir290* SE (Whyte et al., 2013) (**Figure 5G, Figure 5-figure supplement 1C**). The remaining Oct4-bound transcribed enhancers showed no changes in eRNA synthesis and chromatin accessibility upon Oct4 depletion (**Figure 5-figure supplement 1D**). We hereafter refer the three groups of Oct4-bound transcribed enhancers as Oct4-sensitive, Oct4-insensitive and Oct4-independent enhancers (**Figure 5C**). In conclusion, these results indicate that Oct4 binding to putative enhancers does not directly correlate with chromatin accessibility.

### Sox2 may contribute to retained enhancer accessibility upon Oct4 depletion

We then asked whether Sox2 may contribute to the delayed decrease in chromatin accessibility observed at Oct4-sensitive enhancers (**Figure 5D**). We performed ChIP-seq of Sox2 over the same time course (Table S6). At Oct4-sensitive enhancers, Sox2 remained bound from 0 to 6 hours and started to decrease after 9 hours of treatment (**Figure 6A-B**). At Oct4-insensitive and Oct4-independent enhancers, Sox2 occupancy was stable over the entire time course (**Figure 6A-B, Figure 6-figure supplement 1A-B**). Moreover, we analyzed published Oct4, Sox2 and Nanog ChIP-seq data at 0 and 24 hours after DOX treatment (King and Klose, 2017). Oct4-sensitive enhancers showed a strong loss of all three TFs at 24 hours, whereas Sox2 occupancy was unchanged and Nanog occupancy increased at 24 hours at Oct4-insensitive enhancers and Oct4-independent enhancers (**Figure 6-figure supplement 1C-E**).

**Figure 6.**
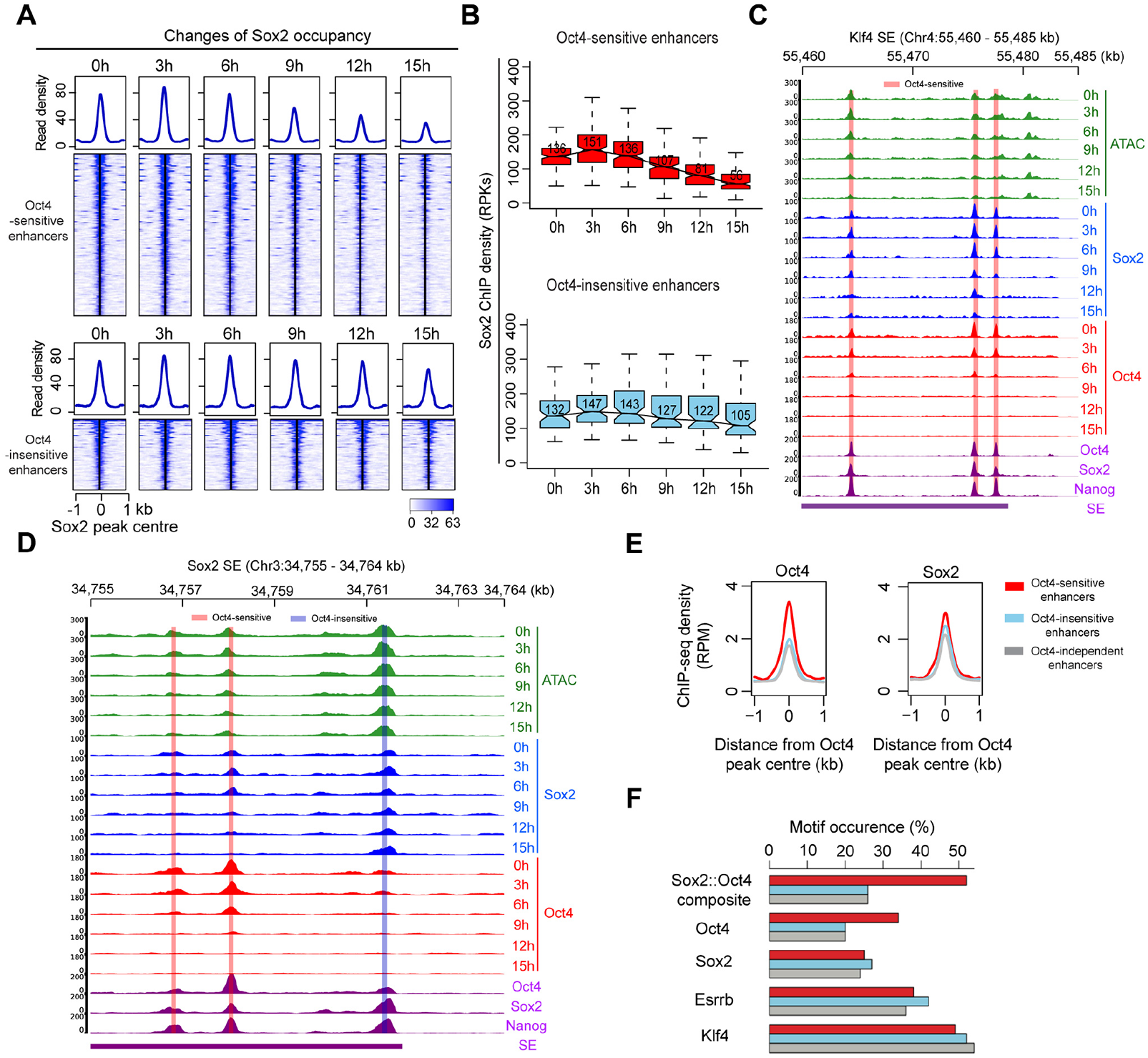
Sox2 may contribute to retained enhancer accessibility upon Oct4 depletion. **(A)** Heatmap showing changes of Sox2 occupancy at Oct4-sensitive and Oct4-insensitive enhancers over the entire time course of DOX treatment. Normalized read densities were presented, peaks were ranked by Sox2 read densities. **(B)** Same as **(A)**, but using boxplots to depict quantification of Sox2 occupancy changes at Oct4-sensitive and Oct4-insensitive enhancers. Y axis represents Sox2 ChIP-seq density in reads per kilobases (RPKs). Black bars represent the median values for each group. Lower and upper boxes are the first and third quartiles, respectively. The ends of the whiskers extend the box by 1.5 times the interquartile range. Outliers are omitted. **(C)** Genome browser view illustrating changes of chromatin accessibility, Sox2 and Oct4 occupancy at *Klf4* SE. Tracks from top to bottom: ATAC-seq coverages (green), ChIP-seq coverages for Sox2 (blue) and Oct4 (red) at 0, 3, 6, 9, 12 and 15 hours; ChIP-seq coverages for Oct4, Sox2 and Nanog (purple) from ZHBTc4 mouse ES cell at 0 hours (King and Klose, 2017); SE annotation (Whyte et al., 2013). Biological replicates were merged for visualization. **(D)** Genome browser view illustrating changes of chromatin accessibility, Sox2 and Oct4 occupancy at *Sox2* SE. Tracks were visualized and ordered in the same way as in (**C**). **(E)** Metagenes analysis of Oct4 and Sox2 occupancy at Oct4-sensitive, -insensitive and – independent enhancers at 0 hours, data were obtained from (King and Klose, 2017). Y axis depicts ChIP-seq coverage density in reads per million (RPM). **(F)** Percentage of motif occurrence at Oct4-sensitive, -insensitive and -independent enhancers for Sox2-Oct4 composite motif and Oct4, Sox2, Esrrb and Klf4 motifs.

These findings are well illustrated at the exemplary genomic regions comprising putative SEs of *Klf4*, *Sox2* and *Mir290* (**Figure 6C-D, Figure 6-figure supplement 1F**). Within these regions, we observed a decrease of chromatin accessibility and Sox2 occupancy at the Oct4-sensitive enhancers after 9 hours of DOX treatment (**Figure 6C-D, Figure 6-figure supplement 1F**), whereas the Oct4-insensitive enhancers remained accessible and occupied by Sox2 (**Figure 6D, Figure 6-figure supplement 1F**). Taken together, these findings suggest that Sox2 is involved in temporarily keeping chromatin open during the first 6 hours of DOX treatment for Oct4-sensitive enhancers.

### Oct4 may cooperate with Sox2 to render enhancers accessible

To further characterize the differences between Oct4-sensitive and Oct4-insensitive enhancers, we analyzed publicly available ChIP-seq data. Oct4 and Sox2 co-localize in both enhancer groups, with 95% and 86% of Oct4-sensitive and Oct4-insensitive enhancers, respectively, overlapping with Sox2 peaks (**Figure 6-figure supplement 1G**). In metagene plots, Oct4-sensitive enhancers showed ∼1.5-fold enrichment of Oct4 occupancy compared to Oct4-insensitive enhancers (**Figure 6E**), whereas Sox2, Nanog, Klf4 and Esrrb were only slightly enriched if at all (**Figure 6E, Figure 6-figure supplement 1H**). Oct4-sensitive enhancers also displayed higher levels of H3K27ac, whereas H3K4me1 showed similar levels (**Figure 6-figure supplement 1H**). Oct4-independent enhancers showed lower signals for pluripotency TFs and histone modifications, in line with the observed lower transcriptional activity (**Figure 6E, Figure 6-figure supplement 1H**). According to genomic region enrichment analysis (McLean et al., 2010), Oct4-sensitive enhancers were enriched for stem cell population maintenance (**Figure 6-figure supplement 1I**) and Oct4-insensitive enhancers for neural differentiation and development (**Figure 6-figure supplement 1J**). Enhancers of both types may target the same nearest active gene (**Figure 6D, Figure 6-figure supplement 1K**).

To investigate whether a specific binding motif may be related to the enrichment of Oct4 occupancy in Oct4-sensitive enhancers, we performed motif analysis. Our results showed a strong enrichment for both Oct4 and the canonical composite DNA motifs at Oct4-sensitive enhancers only (**Figure 6F**), whereas no difference was found for other TFs. These findings suggest that Oct4 influences chromatin accessibility preferentially when the Sox2-Oct4 composite motif is present in DNA.

### Sox2 maintains chromatin accessibility in the absence of eRNA synthesis

To gain insights into the change of chromatin accessibility at Oct4-bound sites without detected eRNA synthesis, we analyzed the 12,710 Oct4-bound non-transcribed enhancers (**Figure 3A-B**). PCA analysis showed similar transitions for these enhancers over the time course as observed for Oct4-bound transcribed enhancers (**Figure 7-figure supplement 1A**). After 15 hours of DOX treatment, we detected 4,985 enhancers with significantly reduced chromatin accessibility (**Figure 7-figure supplement 1B-D**, adjusted P-value = 0.01). We refer to them as Oct4-sensitive enhancers and the remaining ones as Oct4-insensitive enhancers (**Figure 7A**). For both enhancer groups, Oct4 occupancy decreased already after 3 hours of DOX treatment (**Figure 7B**), and for Oct4-sensitive enhancers the Oct4 occupancy decrease preceded the decrease in chromatin accessibility (**Figure 7B, top**).

We then investigated whether Sox2 may keep chromatin accessible also at Oct4-bound non-transcribed enhancers. Analysis of the Sox2 binding kinetics revealed a decrease of Sox2 binding at Oct4-sensitive enhancers after 9 hours of DOX treatment (**Figure 7C-D**). Oct4-insensitive enhancers showed only a slight decrease of Sox2 binding at 15 hours. Moreover, we analyzed published Oct4, Sox2 and Nanog ChIP-seq data at 0 and 24 hours after DOX treatment (King and Klose, 2017). Oct4-sensitive enhancers showed a strong loss of all three TFs at 24 hours, whereas at Oct4-insensitive enhancers Sox2 and Nanog occupancies were essentially unchanged (**Figure 7-figure supplement 1E**). This is consistent with our earlier observations at Oct4-bound transcribed enhancers (**Figure 6-figure supplement 1C-E**). Furthermore, metagene and motif analysis revealed an enrichment of Oct4 occupancy and the composite DNA motif at Oct4-sensitive enhancers (**Figure 7-figure supplement 1F-G**). Occupancy of other pluripotency factors and associated histone modifications also revealed a similar pattern as observed for Oct4-bound transcribed enhancers (**Figure 7-figure supplement 1F**). Together, these findings suggest that Sox2 can maintain chromatin accessibility at enhancers in the absence of eRNA synthesis during the first 6 hours of DOX treatment.

**Figure 7.**
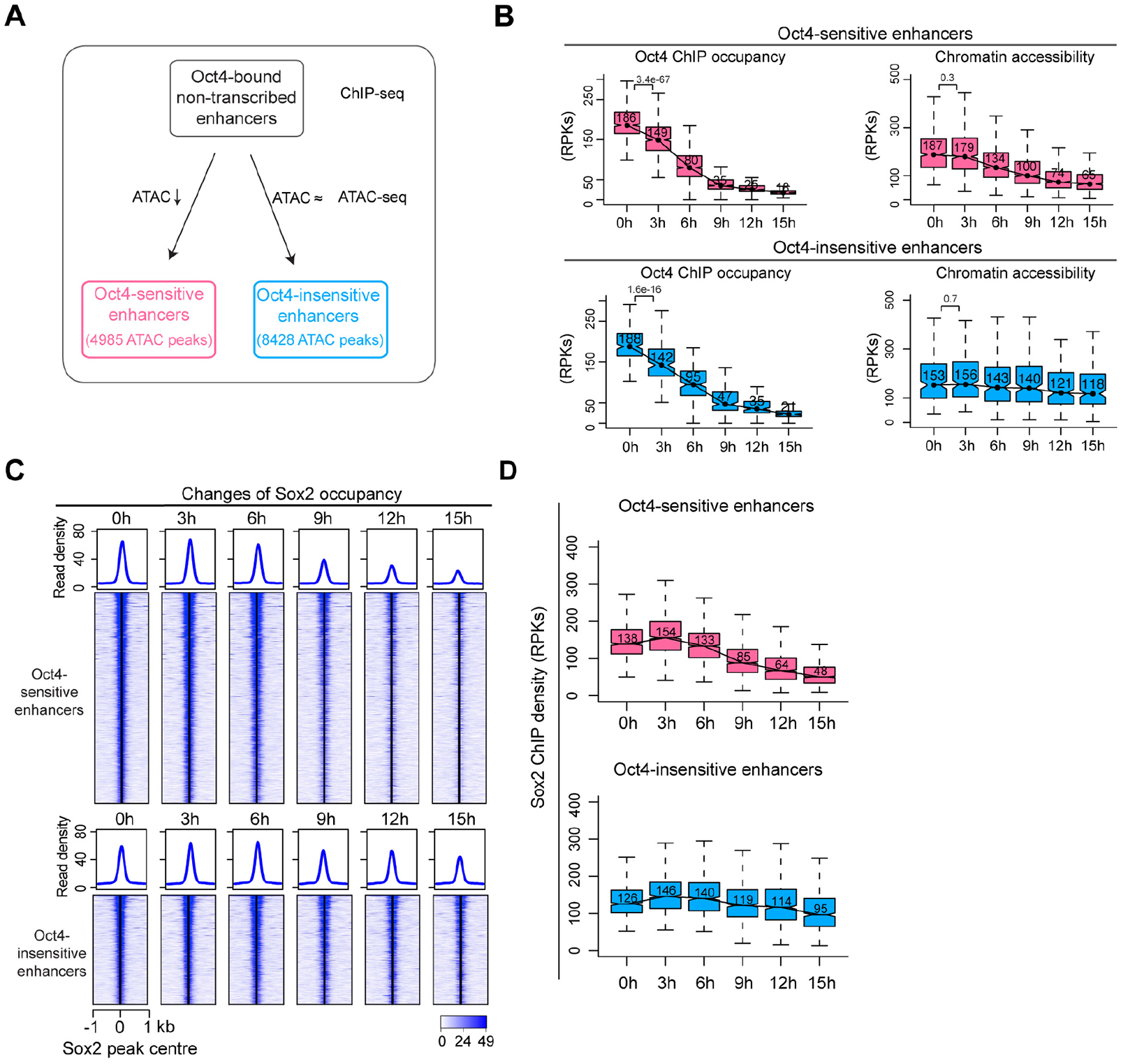
Sox2 maintains chromatin accessibility in the absence of eRNA synthesis. **(A)** Diagram indicating classification of Oct4-sensititive and -insensitive enhancers by changes of chromatin accessibility at Oct4-bound non-transcribed enhancers. **(B)** Boxplots illustrating changes of Oct4 occupancy and chromatin accessibility at Oct4-sensitive and -insensitive enhancers. P-values were calculated by Wilcoxon rank sum test. Y axis represents read counts per kilobases (RPKs). Black bars represent the median values for each group. Lower and upper boxes are the first and third quartiles, respectively. The ends of the whiskers extend the box by 1.5 times the interquartile range. Outliers are omitted. **(C)** Heatmap showing changes of Sox2 occupancy at Oct4-sensitive and Oct4-insensitive enhancers over the entire time course of DOX treatment. Normalized read densities were presented, peaks were ranked by Sox2 read densities. **(D)** Same as (**C**), but using boxplots to depict quantification of Sox2 occupancy changes at Oct4-sensitive and Oct4-insensitive enhancers. Y axis represents Sox2 ChIP-seq density in reads per kilobases (RPKs).

## Discussion

Oct4 and Sox2 are both required for the maintenance of pluripotency, but their individual functions remain incompletely understood. Here we used rapid depletion of Oct4 from mouse ESCs and several genomic approaches to analyze the contributions of Oct4 to the maintenance of pluripotency. Our results show that loss of Oct4 from enhancers goes along with a decrease in mRNA synthesis from putative Oct4 target genes that are involved in maintaining pluripotency. We also observe that Oct4 binding to enhancers is often related to the synthesis of eRNA but is not directly related to enhancer accessibility. Instead, Sox2 can retain enhancer accessibility for some time after Oct4 depletion. This does not depend on eRNA synthesis, indicating that it is not the primary function of eRNA synthesis to keep enhancers accessible.

Our results suggest that Oct4 mainly acts as an activator that stimulates transcription of pluripotency enhancers and their target genes, whereas Sox2 acts as a factor that renders chromatin accessible. Upon Oct4 depletion, a group of enhancers is inactivated in two steps, by a loss of enhancer transcription followed by a decrease in chromatin accessibility. It has been suggested that the activation of enhancers also happens in two steps, first by recruitment of factors that render chromatin accessible, and second by recruitment of factors that induce enhancer transcription (Li et al., 2016). Based on our results and available literature we therefore propose that the activation and inactivation of enhancers both occur in a two-step process with distinct factors dominating each step but also often cooperating.

Consistent with our genome-wide *in vivo* analysis, structural studies found that Sox2 alone can bind a nucleosome and displace part of the nucleosomal DNA from the octamer surface (Dodonova et al., 2020), and that Oct4 and Sox2 can co-occupy a nucleosome, contacting each other (Michael et al., 2020). It was also found that Oct4 and Sox2 bind cooperatively to a composite DNA motif *in vitro* (Chang et al., 2017) and that Sox2 can increase the specificity for Oct4 binding i*n vivo* (Merino et al., 2014; Michael et al., 2020). Also supporting our findings, single molecule imaging showed that Sox2 assists Oct4 binding in live ESCs (Chen et al., 2014).

In conclusion, our data indicate that the primary function of Oct4 is to control enhancer activity rather than accessibility. This is further supported by published results on the functions of Oct4 and Sox2 during reprogramming of somatic cells to induced pluripotent stem cells (iPSC) (Chen et al., 2016; Chronis et al., 2017; Li et al., 2017; Soufi et al., 2012; Soufi et al., 2015; Takahashi and Yamanaka, 2006; Velychko et al., 2019). Several lines of evidences indicate that during reprogramming Sox2 acts as a driver of chromatin opening, whereas Oct4 is an accessory factor (Li et al., 2017; Malik et al., 2019; Soufi et al., 2015; Velychko et al., 2019), that is not strictly required (An et al., 2019; Velychko et al., 2019). Thus, a general model emerges that Oct4 controls enhancer activity, whereas Sox2 governs enhancer accessibility, although both factors often cooperate and engage in functional interactions with other factors.

## Materials and methods

### Employed cell lines and doxycycline treatment

Mouse embryonic stem cells (mESCs) harbouring a doxycycline repressible *Pou5f1* transgene (Niwa et al., 2000) were propagated on gelatin-coated plates in equal parts DMEM-F12 (Life Technology, 21331-020) and Neuralbasal (Life Technology, 21103-049) Medium supplemented with 2% Fetal Bovin Serum (Sigma Aldrich, G1393-100 ml), 2% Knockout Serum Replacement Medium (Gibco, 10828-028), 0.04 µg/ml Leukemia Inhibitory Factor (prepared in-house), Pennicilin/Streptomycin (Sigma Aldrich, P4333-100ml), 0.1 mM β-Mercaptoethanol (Gibco, 31350-010), 0.5x B27 supplement (Life Technology, 12587-010), 0.5x N2 supplement (Gibco, AM9759, 3 µM CHIR99021 (Cayman Chemicals, 13122), 1 µM PD0325901 (Biomol, 103034-25). Cells were passaged using Accutase (Sigma Aldrich, 6964-100ml), *Pou5f1* expression was abolished by treatment with 1µg/ml doxycycline (Sigma Aldrich D9891-1G) for 3 hours, 6 hours, 9 hours, 12 hours, 15 hours (verified by western blot) and 24 hours (verified by immunofluoresence and western blot).

### Sample preparation and Western blotting

ZHBTc4 cells were washed with PBS and harvested using Accutase (Sigma Aldrich, 6964-100 ml) at the given time points of loss-of-Oct4. Cells were centrifuged for 5 minutes at 1400 rpm, supernatant was aspirated and cell pellet re-suspended as single cell suspension in cell culture medium. For whole cell lysate, cell pellets were weighed. Cell pellets were re-suspended in 4x LDS buffer (prepared in-house) based on weight. Equal volume for each sample was loaded on to SDS-PAGE gels. For chromatin samples, cell number was determined using counting chamber. 2×10^7^ cells were cross-linked with final concentration of 1% formaldehyde AppliChem, A0877,0250) for 8 minutes and quenched for 5 minutes with 125 mM Glycine Sigma Aldrich, G8898-1KG). Cross-linked cells were centrifuged for 5 minutes at 1350 x g and washed twice with 1 ml of cold PBS. Cells were either stored at -80°C or directly processed for chromatin extraction. For whole cell lysate samples equal volume for each sample was loaded on to SDS-PAGE gels. Chromatin samples were loaded equally based on DNA concentrations. Blots were probed for Oct4 (Santa Cruz, sc-5279), Sox2 (Santa Cruz, sc17320) Nanog (Bethyl Laboratories, A300-397A), Histone 3 (H3) for chromatin samples (Abcam ab1791), Tubulin for whole cell lysate samples (Sigma Aldrich, T6199) overnight at 4°C. Next day, blots were washed and probed with secondary antibodies anti-mouse-HRP (Jackson Labs, 115-035-044), anti-goat-HRP (R&D systems, HAF019), anti-rabbit-HRP (GE healthcare, NA934) at room temperature for 2 hours. Blots were exposed to film using ECL (GE healthcare, RPN2232).

### Immunofluorescence

ZHBTc4 cells were treated for 0 and 24 hours. Next, cells were cross-linked with 4% paraformaldehyde (Sigma Aldrich, D6148-500G) for 30 minutes. Formaldehyde was quenched with 50 mM Glycine (Sigma Aldrich, G8898-1KG) for 15 minutes. Cells were stained for Oct4 (Santa Cruz sc-5279) or Sox2 (Santa Cruz, sc17320) overnight at 4°C. The following day samples were incubated with anti-goat-alexa-488 (Thermoscientific, A11078) or anti-goat-alexa-568 (Thermoscientific, A11061) and Hoechst (Sigma Aldrich, H6024).

### TT-seq

TT-seq was performed as described (Schwalb et al., 2016) with minor alterations. In brief, two biological replicates at the aforementioned timepoints were produced for TT-seq. 1*10^8 cells were labeled with 500 µM 4-thiouridine (4sU, Carbosynth, 13957-31-8) for 5min. Cells were harvested and lysed using TRIzol (Ambion, 1559018) and stored at -80°C. Prior to RNA isolation, RNA spike-ins were added at 5 ng per 1*10^8 cells. Details regarding the used spike-in sequences and generation of the spike-in mix can be found in (Wachutka et al., 2019). Total RNA was isolated using TRIzol (Ambion, 1559018) according to manufactures instructions, and subsequentially fragmented to 1500-5000 bp using Covaris S220 Ultrasonicator. Nascent RNA was purified as described (Schwalb et al., 2016) with minor modifications. Following purification using streptavidin pulldown, the collected RNA was purified using RNeasy micro Kit (Qiagen, 74004), as well as DNase treatment (Qiagen, 79254). Sequencing libraries were produced using NuGen Ovation Universal RNA-seq System (Nugen, 0343). Size selected libraries were analyzed on a Fragment Analyzer before sequencing on a Illumina NEXTseq 550.

### ATAC-seq

ATAC-seq was performed as described (Buenrostro et al., 2013) with a few alterations. ZHBTc4 cells were harvested using Accutase (Sigma Aldrich, 6964-100 ml) at 0 hours, 3 hours, 6 hours, 9 hours, 12 hours, 15 hours. Nuclear isolation of 5×10^4 cells was followed by treatment with Nextera Tn5 enzyme (Illumina, 20034198) for 45 minutes at 37°C. PCR amplification of the samples was performed using Nextera primers 1 and 2 and NEBNext High fidelity master mix (NEB, M0541S) for 12 cycles as determined by KAPA Real-Time Library Amplification Kit (Peqlab, KK2701). Libraries were purified over Machnery-Nagel PCR spin column (Macherey Nagel, 740609.50S) and AMPure XP beads (Beckman Coulter, A63881) in a 1:1.8 ratio. Sequencing of libraries was performed on a Illumina NEXTseq 550.

### Sox2 ChIP-seq

Cross-linked pellets (as described in sample preparation for western blot) were thawed on ice. Protease inhibitor (Roche, 4693124001) was added to all buffers. Pellets were resuspended in lysis buffer 1 (50 mM Hepes-KOH pH7.5, 140 mM NaCl, 1 mM EDTA pH8, 10% glycerol, 0.5% IGEPAL CA630, 0.25% Triton-X100) and lysed for 30 minutes on ice. Samples were pelleted and washed with lysis buffer 2 (10 mM Tris-HCl pH8, 200 mM NaCl, 1 mM EDTA, 0.5M EGTA) for 10 minutes on roller bank at 4 C. Samples were pelleted and re-suspended in SDS sonication buffer (10mM Tris-HCl, 1mM EDTA, 0.5% SDS), incubated on ice for 10 minutes and transferred to TPX sonication tubes (Diagnod, C30010009). Chromatin was sonicated in Diagnod Bioruptor 4 x 15 minutes at 30 seconds ON and 30 seconds OFF, high setting in a cooled water bath. Sheared chromatin was centrifuged for 10 minutes 15000 rpm at 4°C. 25 µl of sample was de-crosslinked overnight at 65°C and distribution of size was checked on 1.4% agarose gel. 3µg of Sox2 antibody (Neuromics, GT15098) was coupled to Dynabeads protein G (Thermo Fisher Scientific, 10009D) for 2 hours at 4°C for each sample. 50µg of chromatin was used for each immunoprecipitation (IP). Chromatin was diluted using ChIP dilution buffer (10mM Tris-HCl pH8, 125mM NaCl, 0.125% Sodium deoxycholate, 1.25% Triton-X100). Antibody-chromatin mix was incubated overnight at 4°C rotating end-over-end. Samples were washed with low salt buffer (20mM Tris-HCl pH8, 150mM NaCL, 2mM EDTA, 0.1% SDS, 1% Triton-X100), twice using high salt buffer (20mM Tris-HCl pH8, 500mM NaCl, 0.1% SDS, 1% Triton-X100), twice using RIPA washing buffer (50mM HEPES-KOH pH7.6, 250mM LiCl, 1mM EDTA, 1% IGEPAL CA630, 0.7% Sodium deoxycholate) and once with TE buffer containing 50mM NaCl. Bound chromatin was eluted using 105µl pre-warmed elution buffer (10mM Tris-HCl pH8, 5mM EDTA, 300mM NaCl, 0.5% SDS) for 15 minutes at 65°C. RNase A (Invitrogen, 1004D was added and the samples were incubated overnight at 65°C. Next day, samples were treated with Proteinase K (AppliChem, A4392,0010) for 2 hours at 55°C. Samples were purified using Machnerey-Nagel PCR spin column (Macherey Nagel, 740609.50S). DNA quantity was done using Qubit 3.0 (Life Technology, Q33126). 25ng of DNA was used to prepare sequencing libraries using NEBNext® Ultra™ DNA Library Prep Kit (NEB, E7370L) according to manufactures manual. Purity and size distribution was analyzed on Fragment Analyzer. Libraries were sequenced on a HiSeq 1500 (Illumina).

### Oct4 ChIP-seq

Cross-linked pellets (as described in sample preparation for western blot) were thawed on ice. Protease inhibitor (Roche, 4693124001) was added to all buffers. A pellet coming from 3 x 10^7^ cells was resuspended in Farnham Lysis buffer (5 mM Pipes pH 8, 85 mM KCl, 0.5 % NP-40) and lysed for 10 minutes on ice. Samples were pelleted for 5 min at 1,700 g at 4 °C. Samples were washed with PBS and pelleted for 5 min at 1,700 g at 4 °C. Samples were re-suspended in 1 mL of SDS sonication buffer (10 mM Tris-HCl 7.5 pH, 1 mM EDTA, 0.4 % SDS), incubated on ice for 10 minutes and transfer to AFA milliTube. Sonication was performed with a S220 Focused-ultrasonicator (Covaris) with the following parameters: duty cycle 5 %, peak incident power 140 W, cycle per burst 200, processing time 840 sec, degassing mode continuous, water run level 8. 25 µl of sample was de-crosslinked overnight at 65°C and distribution of size was checked on 1.4% agarose gel. 40 µg of Oct4 antibody (R&D, AF1759) was coupled to Dynabeads protein G (Thermo Fisher Scientific, 10009D) for 2 hours at room temperature for each sample. 100 µg of chromatin was used for each IP. 100 ng of *Drosophila S2* sheared crosslinked-chromatin (Covaris S200 parameters: duty cycle 5 %, peak incident power 140 W, cycle per burst 200, processing time 1,800 sec, degassing mode continuous, water run level 8) were added to 100 μg of chromatin as spike-ins control. Chromatin was diluted in IP buffer (50 mM Hepes pH 7.9, 150 mM NaCl, 1 mM EDTA, 1 % Triton X-100, 0.1 % Sodium-deoxycholate) to obtain a 0.1 % final concentration of SDS. 1 % of diluted chromatin was kept as input at 4 °C. Antibody-chromatin mix was incubated overnight at 4°C rotating end-over-end. Samples were washed 5 times with IP wash buffer (100 mM Tris HCl pH 7.5, 500 mM LiCl, 1 % NP-40, 1 % Sodium-deoxycholate) and one time with TE buffer (10 mM Tris-HCl pH 8, 1 mM EDTA). Immuno-bound chromatin was eluted at 70 °C for 10 min with elution buffer (0.1M NaHCO3, 1 %SDS) and de-crosslinked overnight at 65 °C. After RNAse A treatment at 37 °C for 1.5 h and proteinase K treatment at 45 °C for 2 h, DNA was extracted with one volume phenol:chloroform:isoamyl alcohol 25:24:1 (Sigma-Aldrich, P2069) and precipitated for 30 min at -80 °C with 200 mM NaCl and 100 % ethanol. Pellet was washed with 70 % ethanol and resuspended in TE buffer. DNA quality and size distribution were checked on Fragment Analyzer. 3 ng of DNA was used for library preparation according to NEBNext® Ultra™ II DNA Library Prep Kit (NEB, E7645S). Purity and size distribution was analyzed on Fragment Analyzer. Size-selected libraries were sequenced on Illumina NEXTseq 550.

### TT-seq data preprocessing

Paired-end 42 bp reads were mapped to the mouse genome assembly mm10 using STAR 2.5.3 (Dobin et al., 2013) with the following parameters: outFilterMismatchNmax 2, outFilterMultimapScoreRange 0 and alignIntronMax 500,000. SAMtools (Li et al., 2009) was then used to remove alignments with MAPQ smaller than 7 (-q 7) and only proper pairs (-f 2) were selected. HTSeq-count (Anders et al., 2015) was used to calculate fragment counts for different features. Further data processing was carried out using the R/Bioconductor environment.

### Transcription unit annotation and classification

Annotation of transcription unit was performed as described (Schwalb et al., 2016) with minor modifications. Briefly, the whole genome was segmented into 200-bp consecutive bins and the midpoint of TT-seq fragments was then used to calculate the coverage for each bin for each sample. A pseudo-count was added to each bin to avoid noisy signals. In order to create a unified annotation independent of a specific time point, all TT-seq samples were combined. The R/Bioconductor package GenoSTAN (Zacher et al., 2017) was then used to learn a two-state hidden Markov model with a PoissonLog-Normal emission distribution in order to segment the genome into “transcribed” and “untranscribed” states. Transcribed regions overlapping at least 20% of their length with GENCODE annotated protein-coding gene or lincRNA and overlapping with an annotated exon were classified as mRNA/lincRNA and the rest was defined as ncRNA. Transcribed regions mapping to exons of the same protein-coding gene or lincRNA were combined to create a consecutive transcription unit. In order to avoid spurious predictions, ATAC-seq data was used to call open chromatin regions (see below) and TUs without their promoter (+/-1kb of transcription start site, TSS) overlapping with an opening chromatin region were removed (**Figure 1-figure supplement 1D**). A minimal expression threshold was optimized based on expression difference between TUs with or without their promoter overlapping an opening chromatin region. This resulted in 26822 TUs originating from an open chromatin region with a minimal RPK of 26.5 (**Figure 1-figure supplement 1E**). In order to overcome low expression or mappability issues, ncRNAs that are only 200bp (1bin) apart were merged. Subsequently, TU start and end sites were refined to single nucleotide precision by finding borders of abrupt coverage increase or decrease between two consecutive segments in the four 200-bp bins located around the initially assigned start and stop sites via fitting a piecewise constant curve to the TT-seq coverage profiles for both replicates using the segmentation method from the R/Bioconductor package (Huber et al., 2006) ncRNAs were then classified into the following four categories according to their respective location relative to protein-coding genes: upstream antisense RNA (uaRNA), convergent RNA (conRNA), antisense RNA (asRNA) and intergenic RNA (incRNA) (Figure 1C). ncRNAs located on the opposite strand of an mRNA were classified as asRNA if the TSS was located > 1 kbp downstream of the sense TSS, as uaRNA if the TSS was located < 1 kbp upstream of the sense TSS, and as conRNA if the TSS was located < 1 kbp downstream of the sense TSS. The remaining ncRNAs were classified as incRNA.

To annotate putative eRNAs we selected asRNAs and incRNAs. Since highly synthesized mRNAs can give rise to spurious and un-continuous downstream transcription signal, we restricted the analysis to a subset of asRNA and incRNA that are located 1kb far away from promoter related RNAs including mRNA, uaRNA, conRNA and defined them as putative eRNA. We then merged them if the putative eRNAs fell within 1kb of each other. eRNAs within 1kb of Oct4 occupied opening regions were defined as Oct4-regulated eRNAs and the corresponding Oct4 peaks were classified as Oct4-bound transcribed enhancers. Genome browser view for coverages and annotations were plotted by software pyGenomeTracks (Lopez-Delisle et al., 2021).

### Differential gene expression analysis

R/Bioconductor package DESeq2 (Love et al., 2014) was used to call differentially expressed mRNAs and eRNAs applying DESeq2’s default size factor normalization. For both cases DESeq2 size factors were calcualted using counts for protein-coding genes. An adjusted P-value of 0.01 was used to identify significantly changed mRNAs or eRNAs by comparing each time point to the 0 hours measurement.

### Principal component analysis

For each replicate, size factor normalized feature counts were obtained and DESeq2’s default variance stabilizing transformation was applied. The DESeq2 plotPCA function was used to generate PCA plots with the default parameters.

### K-means clustering

Size factor normalized feature counts were aggregated for two biological replicates for each time point and then the data matrix was subjected to Z-score transformation before clustering. k-means clustering was then performed using the kmeans function in R.

### ATAC-seq data processing

Paired-end 76 bp reads were obtained for each of the samples and Nextera Transposase adapter sequence was removed using (Martin, 2011). Bowtie2 (Langmead and Salzberg, 2012) was used to align paired-end reads to the mouse genome assembly mm10 with the “--local” and “-- no-discordant” options. SAMtools (Li et al., 2009) was then used to remove alignments with MAPQ smaller than 7 (-q 7) and only proper pairs (-f 2) were selected. Reads mapped to custom blacklist regions and mitochondria were removed. Two replicates were pooled and MACS2 (Zhang et al., 2008) was used to call chromatin opening peaks with options: -f BAMPE -g mm --broad --broad-cutoff 0.05. Peaks for all time point were then merged to create non-overlapping unified peaks. Further data processing was carried out using the R/Bioconductor environment.

For quantitative comparison, HTSeq-count (Anders et al., 2015) was used to calculate fragment counts for the non-overlapping unified peaks and DESeq2 was used to call regions with significantly changed chromatin accessibility. For normalization, ATAC-seq peaks overlapping with promoters of protein-coding genes with unchanged mRNA expression were used to calculate DESeq2 size factors. An adjusted P-value of 0.01 was used to identify significantly changed regions by comparing each time point to the 0 hour measurement.

### ChIP-seq data processing

Paired-end ChIP-seq data processing was done as described for ATAC-seq data. For single-end ChIP-seq data, Bowtie2 (Langmead and Salzberg, 2012) was used for mapping with “--local” option. SAMtools (Li et al., 2009) was then used to remove alignments with MAPQ smaller than 7 (-q 7). All published ChIP-seq data were fully processed by ourselves. Detailed information for all ChIP-seq samples can be found in (**Table S4-6**). For peak calling of published paired-end Oct4, Sox2 and Nanog ChIP-seq data, three replicates were pooled and MACS2 (Zhang et al., 2008) was used to call peaks with options: -f BAMPE -g mm. For our paired-end Oct4, Sox2 ChIP-seq data, peaks were called by MACS2 with the same options for each individual replicate. For H3K4me1 peak used in (**Figure 3-figure supplement 1E**), peaks were called by MACS2 with options: -g mm.

For our Oct4 ChIP-seq data, data was normalized using added *D. melanogaster* RNA spike-ins. Normalization factors were obtained by dividing the total *D. melanogaster* spike-ins read counts for each sample by the total spike-ins read counts of the sample with the lowest spike-ins read counts. For our Sox2 ChIP-seq data, data was normalized using total number of uniquely mapped reads. ChIP-seq coverages were divided by the respective normalization factors. HTSeq-count (Anders et al., 2015) was used to calculate fragment counts for peak features. Reads counts were divided by the respective normalization factors.

To avoid noisy signal, we only used Sox2 peaks that were detected by both replicates (consensus peaks) at 0 hours samples for Sox2 occupancy analysis. We overlapped Sox2 consensus peaks with Oct4-sensitive, -insensitive and -independent enhancers and kept the overlapped peaks for the analysis in (**Figure 6A-B**, **7C-D**). For Oct4-bound transcribed enhancers this resulted in 154, 67 and 111 Sox2 consensus peaks that is overlapped with Oct4-sensitive, -insensitive and -independent enhancers in (**Figure 6A-B**). For Oct4-bound non-transcribed enhancers this resulted in 1010 and 478 Sox2 consensus peaks that is overlapped with Oct4-sensitive and -insensitive enhancers in (**Figure 7C-D**). For Oct4 occupancy analysis in (**Figure 7B**), we used the same strategy and this resulted in 844 and 187 Oct4 consensus peaks for Oct4-sensitive and -insensitive enhancers used in (**Figure 7B**).

### GO enrichment analysis

The gene ontology enrichment analysis for differentially expressed mRNAs was performed by DAVID Bioinformatics Resources (Huang da et al., 2009). Genomic regions enrichment analysis was performed by GREAT (McLean et al., 2010).

### Transcription factor binding motif analysis

DNA sequences +/-500bp around Oct4 ChIP-seq peak summit were extracted and FIMO (Grant et al., 2011) was used to find individual motif occurrences. DNA motifs of Oct4, Sox2, Sox2::Oct4 composite, Klf4 and Esrrb were download from JASPAR database (Fornes et al., 2020). Percent of motif occurrence was calculated by counting how many sequences contained the queried motifs in the total of subjected sequences.

### Super enhancer annotation in mESC

Previously, 231 large enhancer domains were identified to be super-enhancers from 8794 sites that are co-occupied by Oct4, Sox2 and Nanog in mESC (Whyte et al., 2013). We downloaded the annotation from the original publication and used UCSC LiftOver (Hinrichs et al., 2006) to convert the coordinates from mouse mm9 genome assembly to mm10.

## Acknowledgements

We thank Kerstin Maier and Petra Rus for sequencing. LX was supported by the International Max Planck Research School for Genome Science, part of the Göttingen Graduate School for Neurosciences, Biophysics, and Molecular Biosciences. LC was supported by EMBO Long-Term Postdoctoral Fellowship (ALTF-1261-2014). P.C. was supported by the Deutsche Forschungsgemeinschaft (SFB860, SPP1935, EXC 2067/1-390729940) and the European Research Council (advanced investigator grant CHROMATRANS, grant agreement no. 882357). EAT and KA were supported by funds of the Max Planck Society.

## Lead contact

Further information and requests should be sent to the lead contact, Patrick Cramer.

## Materials availability

Unique reagents generated in this study are available from the lead contact without restriction.

## Data availability

The sequencing data and processed files and annotations have been submitted to the GEO database (GSE174774).

## Author contributions

The initial design of the study was by EAT, KA and HRS, with input from JC and PC. LX carried out bioinformatics analyses. EAT carried out experiments except that LC performed Oct4 ChIP-seq and JC performed TT-seq, with input from EAT. ML and JC provided assistance with bioinformatics analyses. PC and HRS supervised research. LX, EAT and PC wrote the manuscript with input from LC, ML and HRS.

## Supplemental figures and tables

**Figure 1-figure supplement 1.**
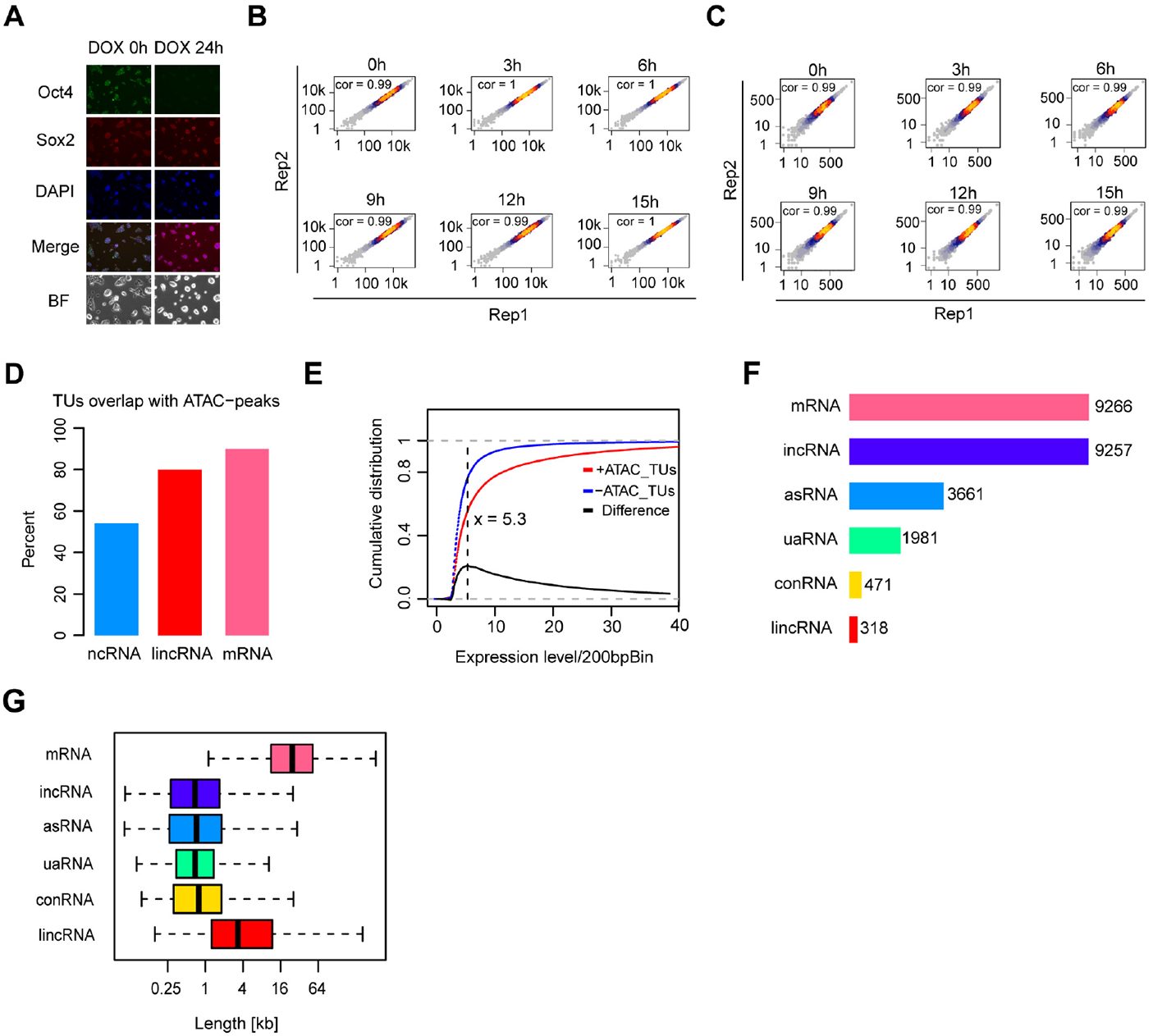
Transcription unit annotation in ZHBTc4 mouse ES cell line, related to Figure 1. **(A)** Immunofluorescence of ZHBTc4 cells after 0 and 24 hours of 1μg/ml DOX treatment. Green: Oct4, red: Sox2, blue: DAPI, BF: Brightfield. **(B)** Scatter plots showing correlation between two biological replicates of TT-seq data at each time point. Normalized TT-seq counts of annotated mRNA (n=9,266) were plotted. Spearman’s rank correlation coefficient was calculated and shown in each plot. **(C)** Scatter plots showing correlation between two biological replicates of ATAC-seq data at each time point. Normalized ATAC-seq counts for Oct4-bound transcribed enhancers (n=2,223) were plotted. Spearman’s rank correlation coefficient was calculated and shown in each plot. **(D)** Bar graph illustrating percentage of overlap of GenoSTAN annotated ncRNAs, lincRNAs and mRNAs with chromatin open regions defined by ATAC-seq peaks. **(E)** Cumulative distribution of expression level of TUs with (red) or without (blue) ATAC-seq peaks overlapping. Black line indicates the difference between distribution of expression level for TUs with or without ATAC-seq peaks overlapping. Dash black line indicates the maximum point for the difference. **(F)** Numbers of different types of annotated TUs. **(G)** Length distribution of different types of annotated TUs.

**Figure 2-figure supplement 1.**
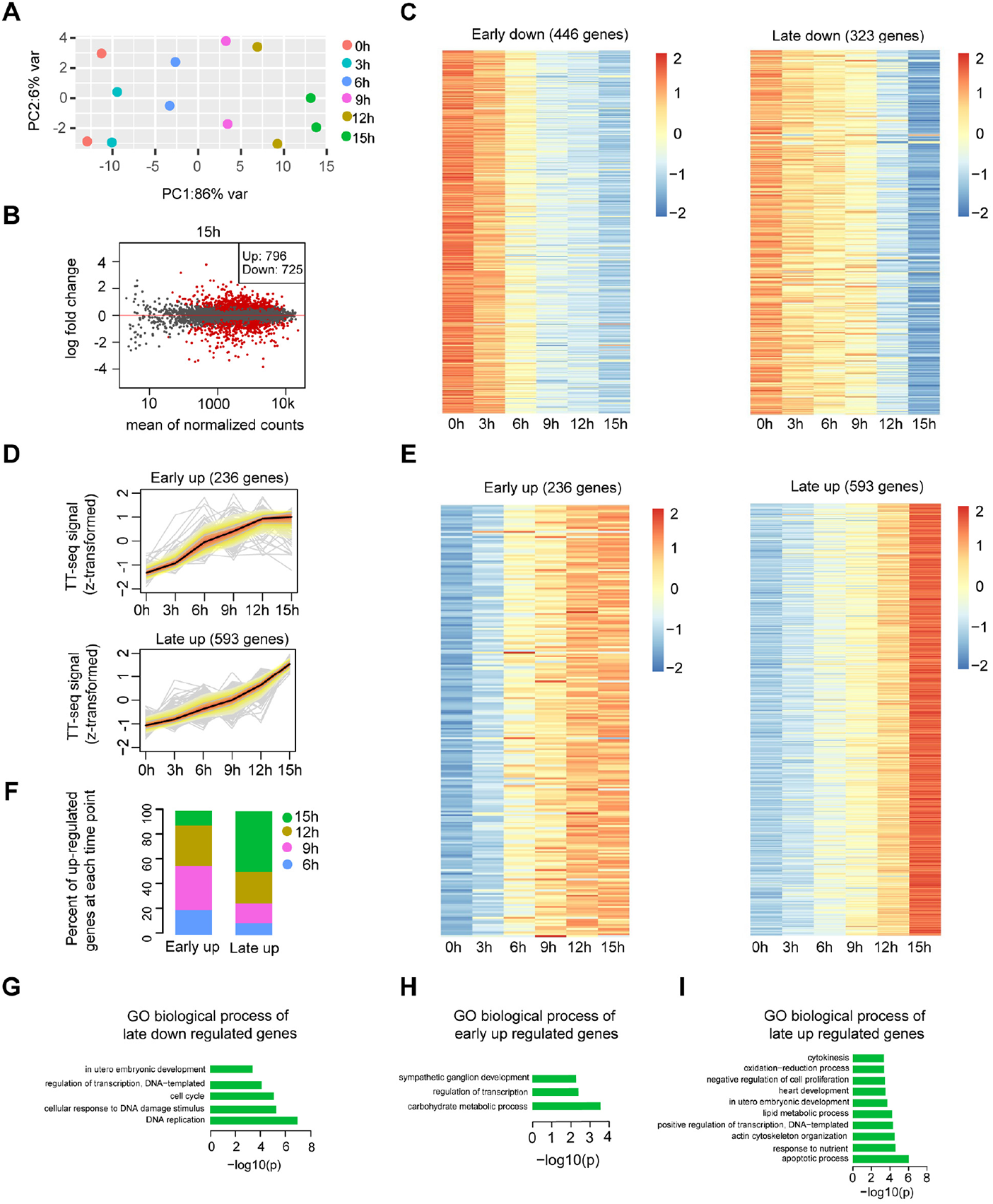
Differential gene expression analysis and clustering, related to Figure 2. **(A)** Principle component analysis of TT-seq annotated mRNA (n=9,226) over time course of DOX treatment (0, 3, 6, 9, 12, 15 hours) for two biological replicates. **(B)** MA plot showing differentially regulated mRNAs detected by DESeq2 after 15 hours of DOX treatment. Statistically significantly regulated mRNAs are depicted in red. **(C)** Heatmap indicating the kinetics of early and late down-regulated genes. **(D)** K-means clustering of up-regulated genes divided into early (top) and late (bottom) up-regulated gene groups. Y axis indicates z-score transformed TT-seq counts. **(E)** Heatmap indicating the kinetics of early and late up-regulated genes. **(F)** Differentially regulated time point composition for early (left) and late (right) up-regulated gene groups. Y axis indicates percentage. **(G-I)** GO biological process enrichment for late down regulated genes (**G**), early up-regulated genes (**H**) and late up-regulated genes (**I**).

**Figure 3-figure supplement 1.**
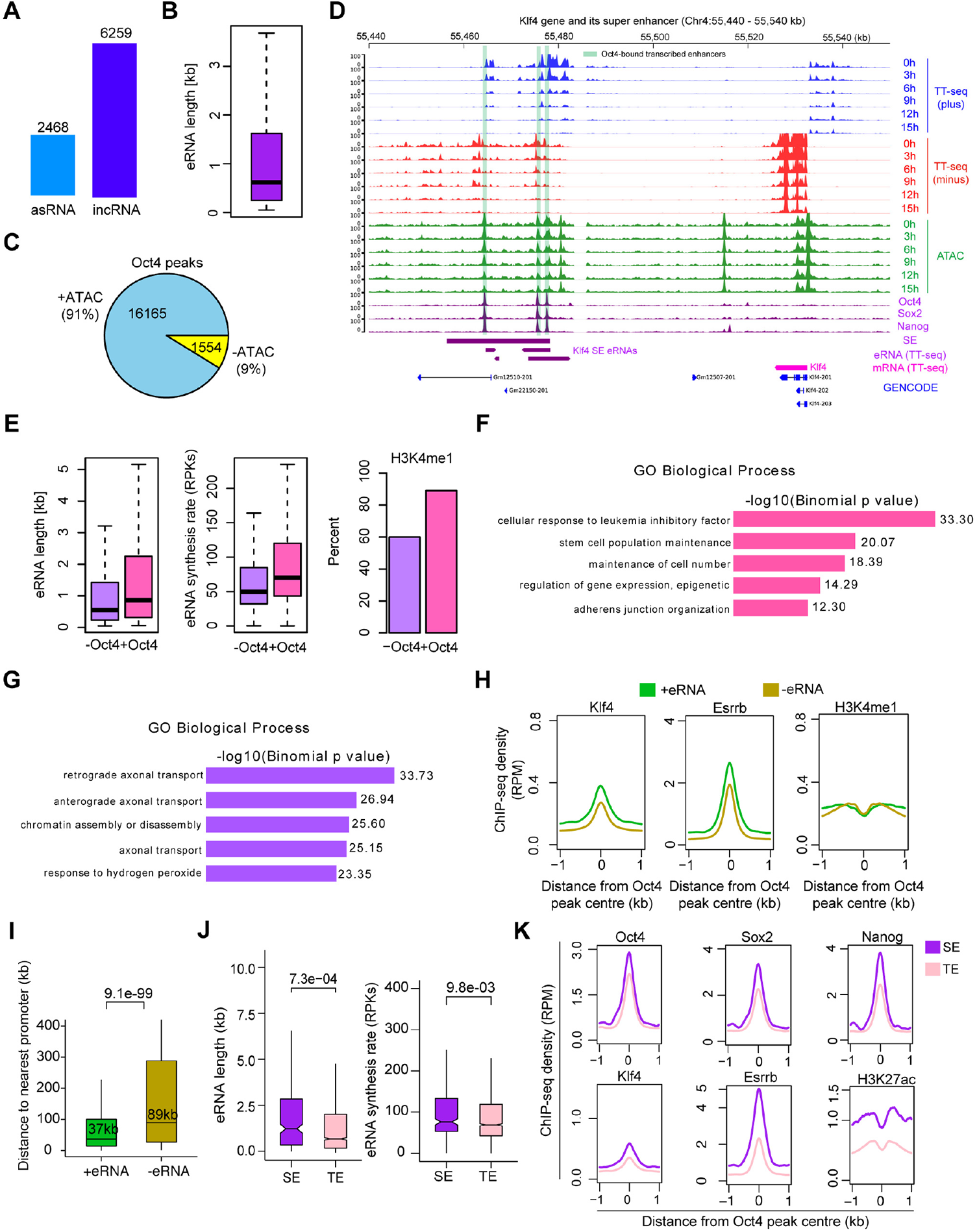
Annotation of eRNA and characterization of Oct4-bound transcribed enhancers in mESC, related to Figure 3. **(A)** Total number of asRNAs and incRNAs classified as putative eRNAs. **(B)** Length distribution of putative eRNAs. **(C)** Percentage of Oct4 binding sites overlapped with chromatin open regions detected by our ATAC-seq data. Oct4 ChIP-seq data was obtained from (King and Klose, 2017). **(D)** Genome browser view of Oct4-bound transcribed enhancers at *Kfl4* SE. Tracks from top to bottom: TT-seq coverages of plus strand (blue), minus strand (red) and ATAC-seq coverages (green) at 0, 3, 6, 9, 12 and 15 hours; ChIP-seq coverages of Oct4, Sox2 and Nanog (purple) from ZHBTc4 mouse ES cell at 0 hours (King and Klose, 2017); SE annotation (Whyte et al., 2013); *Klf4* SE eRNAs annotated by TT-seq (purple arrow); *Klf4* gene (magenta arrow) annotated using TT-seq data; mouse GENCODE annotation for *Klf4* gene (blue). Biological replicates were merged for visualization. **(E)** Length, synthesis and H3K4me1 occupancy of Oct4-regulated eRNAs (n=2,221) versus other eRNAs (n=6,506). ChIP-seq data of H3K4me1 was obtained from (Chronis et al., 2017). (**F-G**) GO biological process enrichment of Oct4-regulated eRNAs (**F**) versus other eRNAs regions (**G**). **(H)** Metagene analysis of Klf4, Esrrb occupancy or histone modification (H3K4me1) at Oct4-bound transcribed enhancers (n=2,231) and Oct4-bound non-transcribed enhancers (n=12,710). ChIP-seq data were obtained from (Chronis et al., 2017). Y axis depicts ChIP-seq coverage density in reads per million (RPM). **(I)** Genomic distance distributions of Oct4-bound transcribed enhancers to their nearest gens versus Oct4-bound non-transcribed enhancers to their nearest genes. P-values were calculated by Wilcoxon rank sum test. **(J)** Length and synthesis (reads per kilobases, RPKs) distributions of SE eRNAs (n=243) versus TE eRNAs (n=1,978). P-value were calculated by Wilcoxon rank sum test. **(K)** Metagene analysis of Oct4, Sox2, Nanog, Klf4, Esrrb occupancy and H3K27ac histone modification at Oct4 occupied TE versus SE. ChIP-seq data of Oct4, Sox2 and Nanog were obtained from (King and Klose, 2017) and Klf4, Esrrb and H3K27ac data from (Chronis et al., 2017). Y axis depicts ChIP-seq coverage density in reads per million (RPM).

**Figure 4-figure supplement 1.**
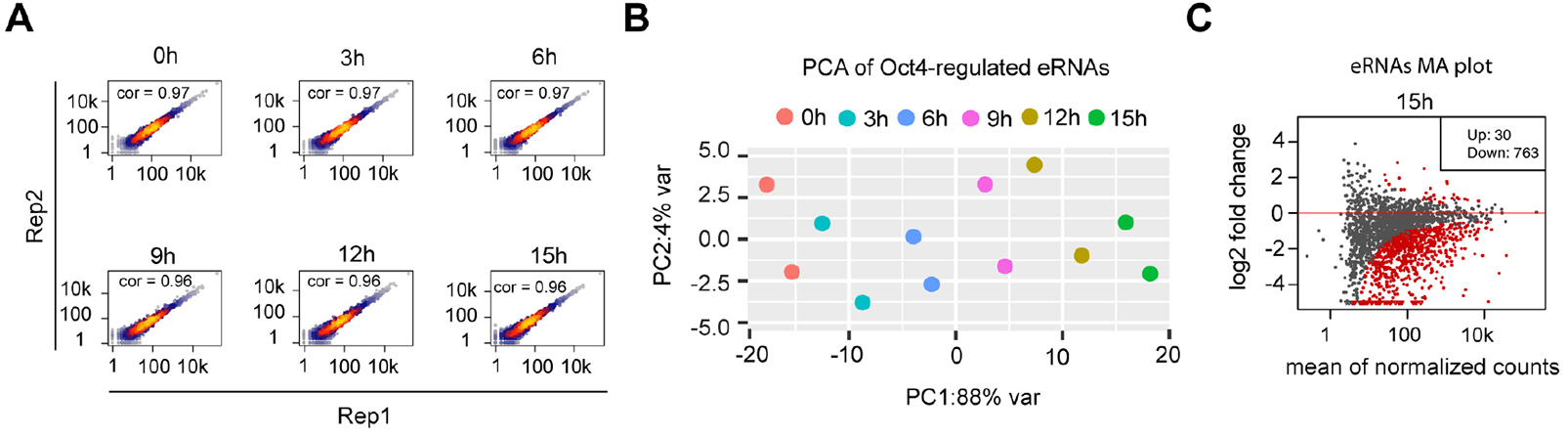
eRNA synthesis changes at Oct4-bound transcribed enhancers, related to Figure 4. **(A)** Scatter plots showing correlation between two biological replicates of TT-seq data for Oct4-regulated eRNAs (n=2,221) for each time point of DOX treatment. Spearman’s rank correlation coefficient was calculated and shown in each plot. **(B)** Principle component analysis of Oct4-regulated eRNA synthesis changes over time-course of DOX treatment (0, 3, 6, 9, 12, 15 hours) for two biological replicates. **(C)** MA plots showing differentially regulated Oct4-regulated eRNAs detected by DESeq2 after 15 hours of DOX treatment. Statistically significantly regulated eRNAs are depicted in red.

**Figure 5-figure supplement 1.**
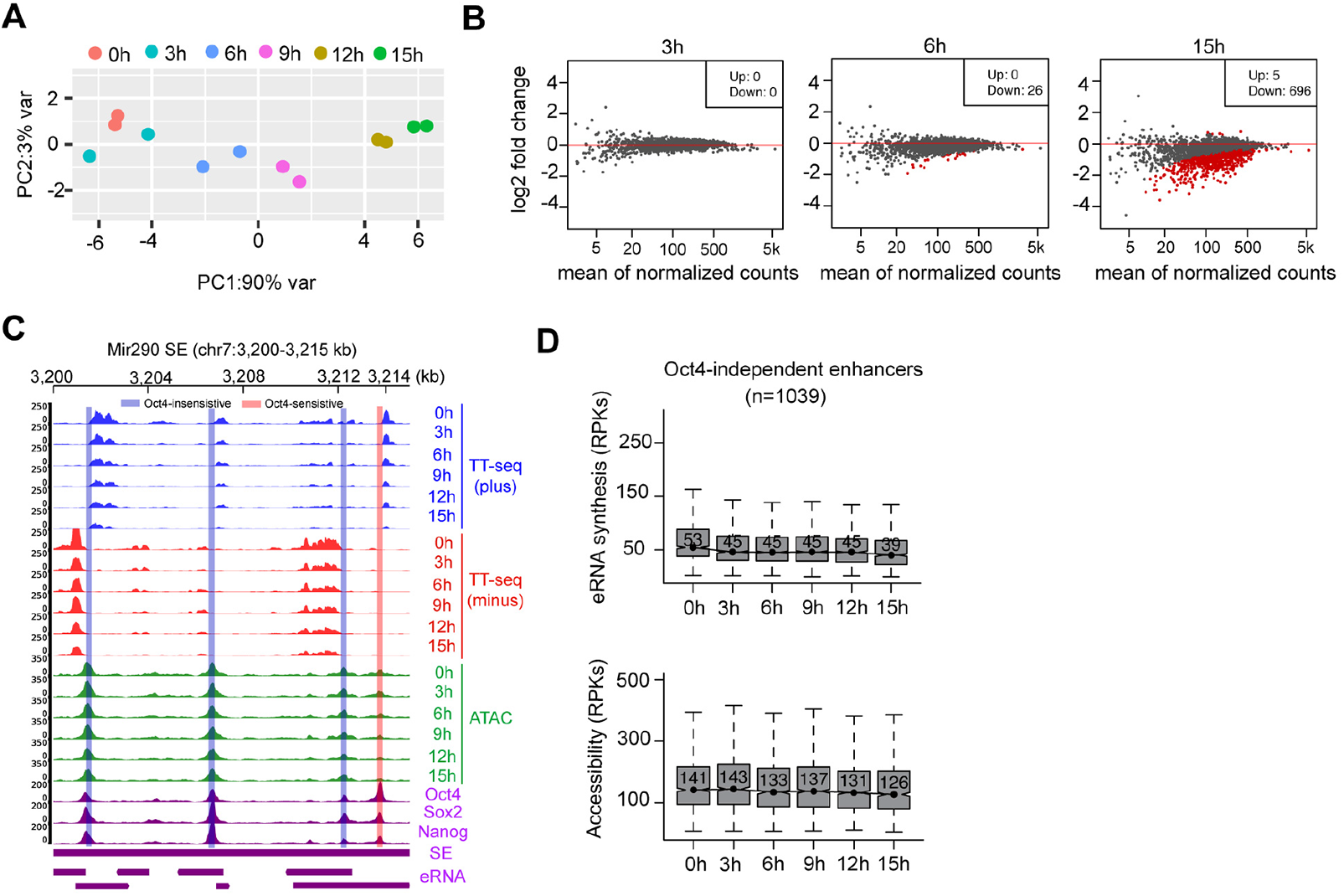
Chromatin accessibility changes at Oct4-bound transcribed enhancers, related to Figure 5. **(A)** Principle component analysis of chromatin accessibility changes at Oct4-bound transcribed enhancers (n=2,223) over time course of DOX treatment (0, 3, 6, 9, 12, 15 hours) for two biological replicates. **(B)** MA plots showing differentially regulated chromatin open regions detected by DESeq2 at 3, 6 and 15 hours DOX treatment for Oct4-bound transcribed enhancers. Statistically significant features are depicted in red. **(C)** Genome browser view for changes of eRNA synthesis and chromatin accessibility at *Mir290* SE including three Oct4-insensitive enhancers and one Oct4-sensitive enhancer. Tracks from top to bottom: TT-seq coverages of plus strand (blue), minus strand (red) and ATAC-seq coverages (green) at 0, 3, 6, 9, 12 and 15 hours; ChIP-seq coverages of Oct4, Sox2 and Nanog (purple) from ZHBTc4 mouse ES cell at 0 hours (King and Klose, 2017); SE annotation (Whyte et al., 2013); *Mir290* SE eRNAs annotated by TT-seq (purple arrow). Biological replicates were merged for visualization. **(D)** Boxplots showing the changes of eRNA synthesis and chromatin accessibility at Oct4-independent enhancers (n=1,039).

**Figure 6-figure supplement 1.**
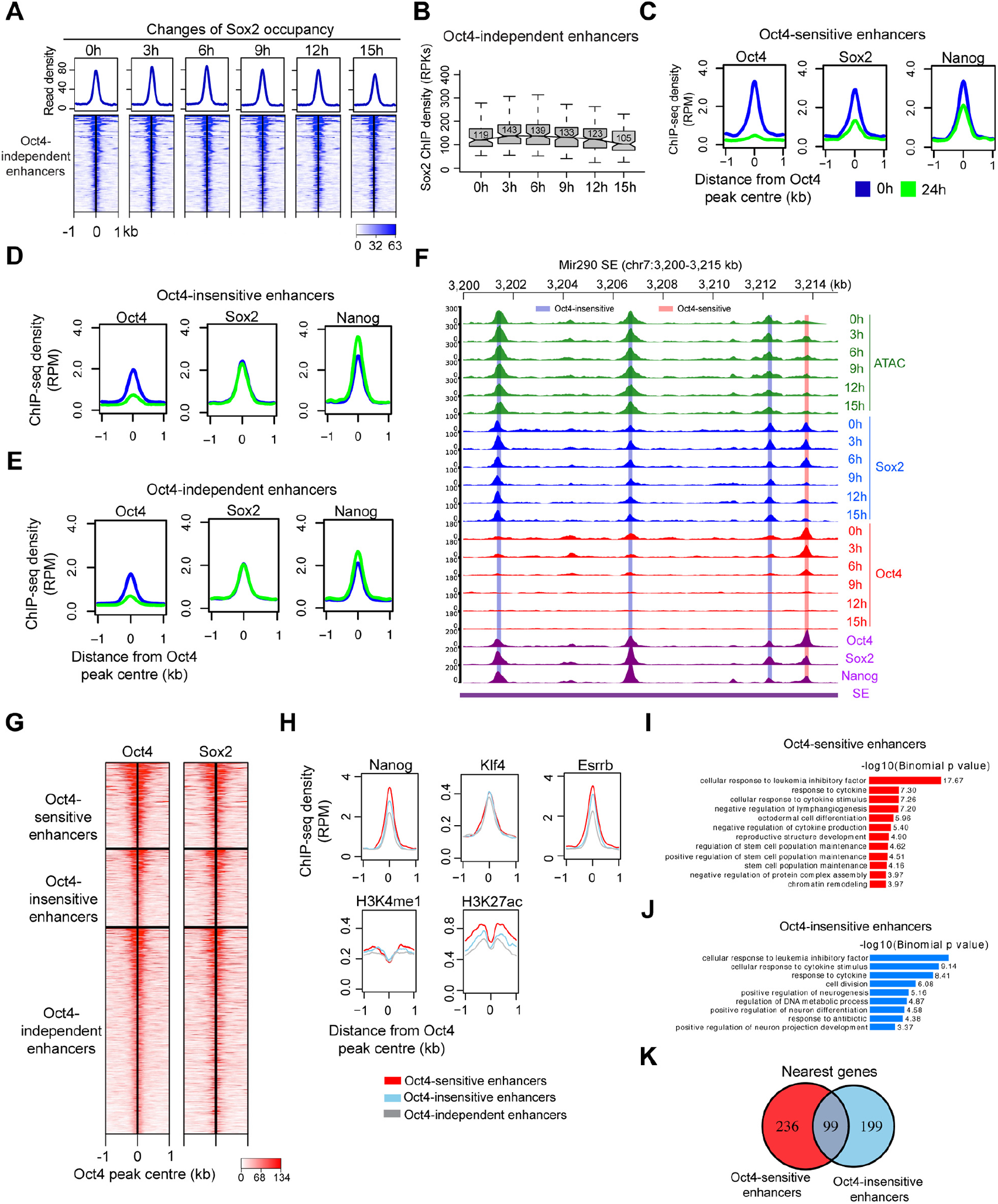
Characterization of Oct4-sensitive, -insensitive and – independent enhancers, related to Figure 6. **(A)** Heatmap showing changes of Sox2 occupancy at Oct4-independent enhancers over the entire time course of DOX treatment. Normalized read densities were presented, peaks were ranked by Sox2 read densities. **(B)** Same as (**A**), but using boxplot to depict quantification of Sox2 occupancy changes at Oct4-independent enhancers. Y axis represents Sox2 ChIP-seq density in reads per kilobases (RPKs). **(C)** Metagenes analysis of Oct4, Sox2 and Nanog occupancy after 0 and 24 hours of DOX treatment at Oct4-sensistive enhancers. ChIP-seq data were obtained from (King and Klose, 2017). Y axis depicts ChIP-seq coverage density in reads per million (RPM). **(D)** Metagenes analysis of Oct4, Sox2 and Nanog occupancy after 0 and 24 hours of DOX treatment at Oct4-insensistive enhancers. **(E)** Metagenes analysis of Oct4, Sox2 and Nanog occupancy after 0 and 24 hours of DOX treatment at Oct4-independent enhancers. **(F)** Genome browser view illustrating changes of chromatin accessibility, Sox2 and Oct4 occupancy at *Mir290* SE. Tracks from top to bottom: ATAC-seq coverages (green), ChIP-seq coverages for Sox2 (blue) and Oct4 (red) at 0, 3, 6, 9, 12 and 15 hours; ChIP-seq coverages for Oct4, Sox2 and Nanog (purple) from ZHBTc4 mouse ES cell at 0 hours (King and Klose, 2017); SE annotation (Whyte et al., 2013). Biological replicates were merged for visualization. **(G)** Heatmap showing co-occupancy of Oct4 and Sox2 at Oct4-sensitive, -insensitive and – independent enhancers at 0 hours. ChIP-seq data were obtained from (King and Klose, 2017). **(H)** Metagenes analysis of Nanog, Klf4, Esrrb occupancy and H3K4me1, H3K27ac histone modifications at Oct4-sensistive, -insensitive and -independent enhancers at 0 hours. Nanog ChIP-seq data was obtained from (King and Klose, 2017), other ChIP-seq data were obtained from (Chronis et al., 2017). **(I)** GO biological process enrichment of Oct4-sensitive enhancers. **(J)** GO biological process enrichment of Oct4-insensitive enhancers. **(K)** Overlap of nearest target genes of Oct4-sensitive and Oct4-insensitive enhancers.

**Figure 7-figure supplement 1.**
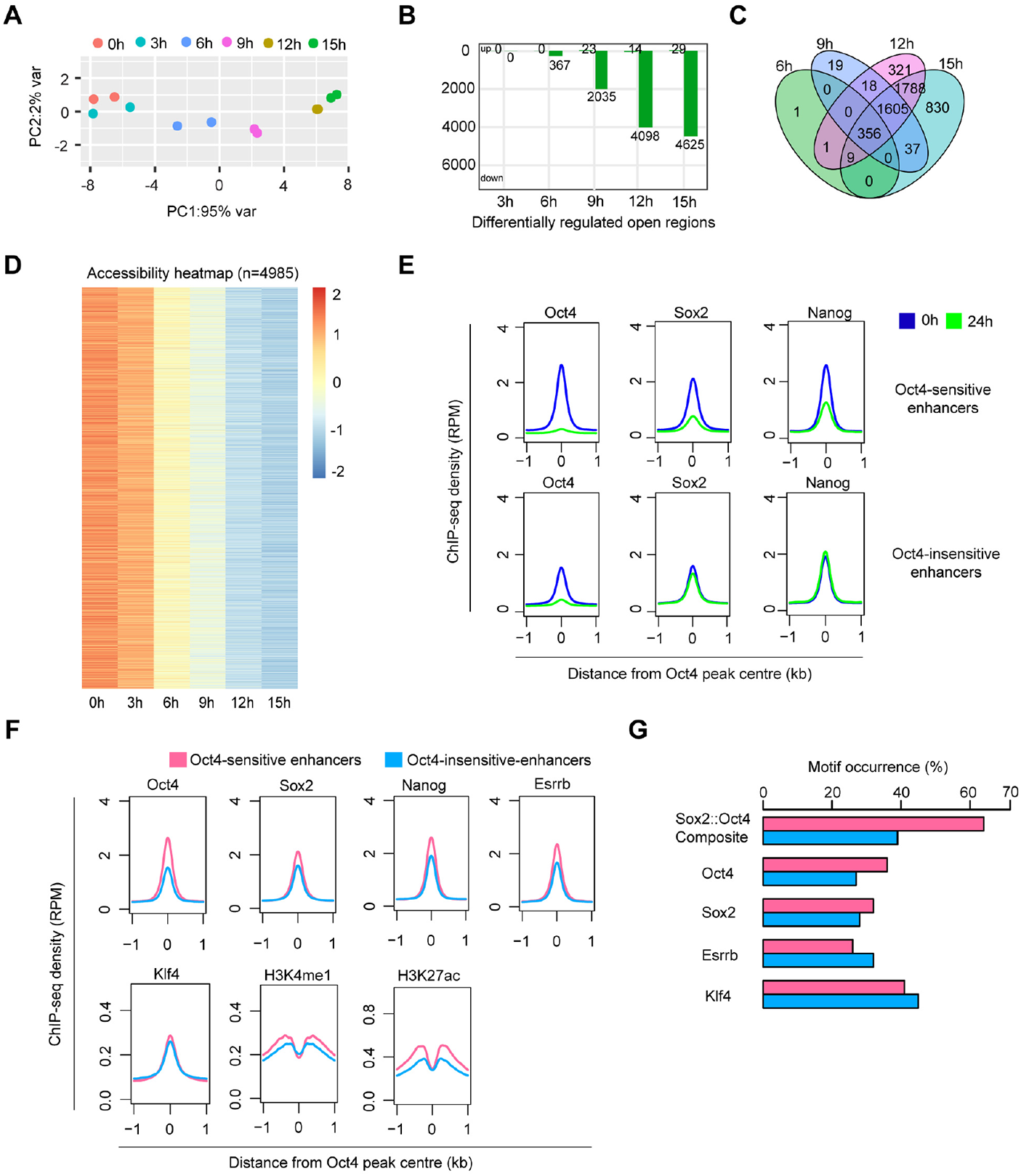
Chromatin accessibility changes at Oct4-bound non-transcribed enhancers, related to Figure 7. **(A)** Principle component analysis of chromatin accessibility changes at Oct4 bound non-transcribed enhancers (n=12,710) over time course of DOX treatment (0, 3, 6, 9, 12, 15 hours) for two biological replicates. **(B)** Number of differentially regulated chromatin open regions detected by DESeq2 at each time point for Oct4 bound non-transcribed enhancers. **(C)** Venn diagram showing overlapping of differentially regulated chromatin open regions from **(B)** at each time points. **(D)** Heatmap visualizing the kinetics of chromatin accessibility changes at differentially regulated chromatin open regions (n=4,985). **(E)** Metagenes analysis of Oct4, Sox2 and Nanog occupancy after 0 and 24 hours of DOX treatment at Oct4-sensitive and -insensitive enhancers. ChIP-seq data of Oct4, Sox2 and Nanog were obtained from (King and Klose, 2017). **(F)** Metagene analysis of Oct4, Sox2, Nanog, Esrrb and Klf4 occupancy and H3K4me1, K3K27ac histone modifications at Oct4-sensitive and insensitive enhancers. ChIP-seq data of Oct4, Sox2 and Nanog were obtained from (King and Klose, 2017), other ChIP-seq data were obtained from (Chronis et al., 2017). Y axis depicts ChIP-seq coverage density in reads per million (RPM). **(G)** Percentage of motif occurrence at Oct4-sensitive and -insensitive enhancers for Sox2-Oct4 composite motif and Oct4, Sox2, Esrrb and Klf4 motifs.

**Table S1.** Sequencing statistics of TT-seq samples generated in this study, related to Figure 1. All samples were sequenced on a NEXTseq 550 sequencing platform in 42bp paired-end mode.

| No. | Hours of DOX treatment | Replicate no. | Sequenced reads | Mapped reads | Duplicates (%) |
| --- | --- | --- | --- | --- | --- |
| 1 | 0h | 1 | 139,767,320 | 115,716,286 | 42.6 |
| 2 |  | 2 | 137,196,527 | 115,935,348 | 31.8 |
| 3 | 3h | 1 | 134,759,731 | 113,952,490 | 31.3 |
| 4 |  | 2 | 140,542,668 | 119,816,299 | 35.4 |
| 5 | 6h | 1 | 139,202,151 | 117,349,804 | 38.5 |
| 6 |  | 2 | 139,904,081 | 118,716,899 | 33.7 |
| 7 | 9h | 1 | 143,258,532 | 120,721,286 | 42.5 |
| 8 |  | 2 | 131,906,425 | 110,880,848 | 35.2 |
| 9 | 12h | 1 | 136,024,166 | 115,730,201 | 38.3 |
| 10 |  | 2 | 136,854,827 | 116,897,968 | 33.4 |
| 11 | 15h | 1 | 141,129,470 | 120,097,702 | 40.9 |
| 12 |  | 2 | 132,219,606 | 111,909,955 | 31.6 |

**Table S2.** Sequencing statistics of ATAC-seq samples generated in this study, related to Figure 1. All samples were sequenced on a NEXTseq 550 sequencing platform in 76bp paired-end mode.

| No. | Hours of DOX treatment | Replicate no. | Sequenced reads | Mapped reads | Duplicates (%) |
| --- | --- | --- | --- | --- | --- |
| 1 | 0h | 1 | 75,879,782 | 60,778,817 | 7.0 |
| 2 |  | 2 | 84,285,910 | 66,234,761 | 7.0 |
| 3 | 3h | 1 | 99,108,163 | 76,192,137 | 7.0 |
| 4 |  | 2 | 95,058,225 | 75,287,164 | 8.0 |
| 5 | 6h | 1 | 100,888,477 | 80,437,745 | 7.0 |
| 6 |  | 2 | 96,527,924 | 77,644,843 | 7.0 |
| 7 | 9h | 1 | 98,029,091 | 76,592,320 | 6.0 |
| 8 |  | 2 | 96,395,215 | 76,365,304 | 7.0 |
| 9 | 12h | 1 | 104,560,309 | 82,912,008 | 7.0 |
| 10 |  | 2 | 91,177,600 | 72,588,902 | 7.0 |
| 11 | 15h | 1 | 93,561,692 | 73,702,559 | 8.0 |
| 12 |  | 2 | 82,330,900 | 65,056,516 | 6.0 |

**Table S3.** Early and late -up and -down regulated gene list, related to Figure 2. See separate excel file.

**Table S4.** List of previously published ChIP-seq datasets used in this study, related to Figure 3 and Figure 3-figure supplement 1.

| No. | Read mode | Data | Hours of DOX treatment | Cell line | Available at |
| --- | --- | --- | --- | --- | --- |
| 1 | Paired-end | Oct4 | 0h | ZHBTc4 | NCBI Gene Expression Omnibus (GSE87822) |
|  |  |  | 24h |  |  |
| 2 | Paired-end | Sox2 | 0h | ZHBTc4 | NCBI Gene Expression Omnibus (GSE87822) |
|  |  |  | 24h |  |  |
| 3 | Paired-end | Nanog | 0h | ZHBTc4 | NCBI Gene Expression Omnibus (GSE87822) |
|  |  |  | 24h |  |  |
| 4 | Single-end | Klf4 |  | C57BL/6J | NCBI Gene Expression Omnibus (GSE 90895) |
| 5 | Single-end | Esrrb |  | C57BL/6J | NCBI Gene Expression Omnibus (GSE 90895) |
| 6 | Single-end | H3K4me1 |  | C57BL/6J | NCBI Gene Expression Omnibus (GSE 90895) |
| 7 | Single-end | H3K4me3 |  | C57BL/6J | NCBI Gene Expression Omnibus (GSE 90895) |
| 8 | Single-end | H3K27ac |  | C57BL/6J | NCBI Gene Expression Omnibus (GSE 90895) |

**Table S5.** Sequencing statistics of Oct4 ChIP-seq samples generated in this study, related to Figure 4 and Figure 7. All samples were sequenced on a NEXTseq 550 sequencing platform in 43bp paired-end mode.

| No. | Hours of DOX treatment | Replicate no. | Sequenced reads | Mapped reads | Duplicates (%) |
| --- | --- | --- | --- | --- | --- |
| 1 | 0h | 1 | 45,489,989 | 37,572,722 | 8.0 |
| 2 |  | 2 | 44,844,975 | 35,827,918 | 9.0 |
| 3 | 3h | 1 | 45,200,637 | 37,381,092 | 7.0 |
| 4 |  | 2 | 40,712,696 | 33,473,643 | 7.0 |
| 5 | 6h | 1 | 35,615,780 | 28,101,180 | 11.0 |
| 6 |  | 2 | 44,105,296 | 34,059,983 | 12.0 |
| 7 | 9h | 1 | 39,889,543 | 32,484,669 | 8.0 |
| 8 |  | 2 | 33,968,970 | 27,705,901 | 8.0 |
| 9 | 12h | 1 | 49,893,843 | 40,471,411 | 9.0 |
| 10 |  | 2 | 40,182,623 | 31,715,336 | 11.0 |
| 11 | 15h | 1 | 54,230,702 | 43,871,081 | 8.0 |
| 12 |  | 2 | 53,405,989 | 38,050,210 | 10.0 |

**Table S6.** Sequencing statistics of Sox2 ChIP-seq samples generated in this study, related to Figure 6-7. All samples were sequenced on a HiSeq 1500 (Illumina) sequencing platform in 76bp paired-end mode.

| No. | Hours of DOX treatment | Replicate no. | Sequenced reads | Mapped reads | Duplicates (%) |
| --- | --- | --- | --- | --- | --- |
| 1 | 0h | 1 | 31,157,808 | 26,708,870 | 8.0 |
| 2 |  | 2 | 30,358,437 | 26,105,044 | 8.0 |
| 3 | 3h | 1 | 29,224,107 | 25,024,914 | 8.0 |
| 4 |  | 2 | 23,143,269 | 19,813,671 | 8.0 |
| 5 | 6h | 1 | 32,774,215 | 28,167,501 | 8.0 |
| 6 |  | 2 | 26,661,227 | 22,987,161 | 7.0 |
| 7 | 9h | 1 | 29,314,133 | 25,170,019 | 9.0 |
| 8 |  | 2 | 25,428,443 | 21,723,313 | 8.0 |
| 9 | 12h | 1 | 26,165,258 | 22,383,270 | 9.0 |
| 10 |  | 2 | 24,258,711 | 20,855,734 | 8.0 |
| 11 | 15h | 1 | 26,728,664 | 22,835,623 | 8.0 |
| 12 |  | 2 | 24,926,639 | 21,369,856 | 9.0 |

